# Isolation and characterization of novel banana rhizosphere bacteria for the control of *Fusarium oxysporum* f. sp. *cubense* TR4

**DOI:** 10.64898/2026.01.29.702532

**Authors:** David-Dan Cohen, Adi Faigenboim, Judith Kraut-Cohen, Marcel Maymon, Stanley Freeman, Shmuel Carmeli, Dror Minz

**Affiliations:** Department of Soil Chemistry, Plant Nutrition and Microbiology, Institute of Soil, Water and Environmental Sciences, Agricultural Research Organization, Volcani Research Center, Beit Dagan, Israel; Department of Agroecology and Plant Health, Robert H. Smith Faculty of Agriculture, Food and Environment, Hebrew University of Jerusalem, Jerusalem, Israel; Institute of Plant Science, Agricultural Research Organization, Volcani Research Center, Beit Dagan, Israel; Department of Plant Pathology and Weed Research, Agricultural Research Organization, Volcani Research Center, Beit Dagan, Israel; School of Chemistry, Raymond & Beverly Sackler Faculty of Exact Sciences, Tel Aviv University, Tel Aviv, Israel

## Abstract

*Fusarium* wilt of banana, caused by *Fusarium oxysporum* f. sp. *cubense* race TR4 (Foc), is one of the most destructive diseases threatening global banana production, particularly the Cavendish cultivar. Conventional control strategies, including chemical treatments and quarantine, remain largely ineffective and unsustainable, underscoring the urgent need for alternative approaches. Biological control using rhizosphere-associated microorganisms offers a promising and environmentally friendly strategy.

In this study, we isolated 436 bacterial strains from the rhizosphere of healthy banana plants and screened them for antifungal activity against Foc. Out of the screened isolates, 93 exhibited significant *in-vitro* inhibitions, and 64 of these were subsequently evaluated in greenhouse assays. We found that 22 strains reduced

Fusarium wilt severity by 45–85% compared to untreated controls. Among them, two isolates, DDC20 and DDC_NEW2, consistently demonstrated strong biocontrol activity. In addition, cell-free culture media (CFCM) and crude extracts inhibited spore germination in fluorescence-based assays, indicating the involvement of secreted antifungal metabolites.

Microscopy and confocal observations of GFP-tagged Foc revealed hyphal abnormalities in the presence of bacterial treatments, including swelling, irregular branching, and distortion, accompanied by excessive sporulation characterized by abundant microconidia, macroconidia, and chlamydospores. Whole-genome sequencing and comparative analyses placed both isolates within the genus *Bacillus*. Genome mining using antiSMASH identified multiple biosynthetic gene clusters encoding known antifungal compounds such as surfactin, fengycin, bacillibactin, and difficidin, as well as putative novel clusters. LC–MS confirmed the presence of surfactin and fengycin in bacterial extracts, supporting the genomic predictions.

Collectively, these findings highlight the potential of DDC20 and DDC_NEW2 (related to *Bacillus* spp.) from the banana rhizosphere as effective biocontrol agents against Foc TR4. This integrated approach, combining phenotypic assays, microscopy, and genome mining, provides a strong foundation for the development of sustainable strategies to manage Fusarium wilt in banana cultivation.

## 1. Introduction

Bananas (*Musa spp.*) are large herbaceous perennials with a branched underground rhizome, abundant roots, and lateral buds [1]. With an annual production exceeding 100 million metric tons and an export value of approximately $5 billion USD [2, 3], bananas rank fourth among global food commodities after rice, wheat, and maize [4, 5].

The global banana industry is increasingly threatened by fungal pathogens [6–8], particularly Panama disease, or Fusarium wilt. This devastating disease is caused by *Fusarium oxysporum* f. sp. *cubense*, a soil-borne pathogen comprising multiple pathogenic races with distinct host specificities. Historically, Race 1 (R1) was responsible for the first major outbreaks of Panama disease, causing the collapse of the Gros Michel banana cultivar but is largely ineffective against Cavendish cultivars. In contrast, *Fusarium oxysporum* f. sp. *cubense* Tropical Race 4 (Foc) represents a highly virulent lineage capable of infecting Cavendish bananas as well as a wide range of additional Musa cultivars.

Since its emergence, Tropical Race 4 has spread rapidly across Asia, Australia, Africa, the Middle East, and more recently South America [9, 10]. Following infection through roots or wounds in the pseudostem, Foc colonizes the vascular system, obstructing xylem vessels and leading to characteristic yellowing of lower leaves, wilting, and ultimately plant death due to severe water imbalance [2, 3]. Cavendish bananas are particularly vulnerable due to clonal propagation through tissue culture, which results in genetic uniformity [7]. This makes entire plantations susceptible to the same pathogens, allowing diseases to spread rapidly [5]. Soilborne pathogens such as Foc are highly persistent: they survive in soil for decades and spread via water, wind, or contaminated tools [11, 12]. Moreover, because early symptoms often mimic abiotic stress, diagnosis is further complicated [13].

Traditional management strategies against Panama disease include field quarantines, soil fumigation, and the application of chemical pesticides [11, 12]. While tissue culture-derived pathogen-free planting material can be used in “clean soil”, these methods are not infallible [14]. The extensive use of chemical pesticides has raised concerns about environmental pollution and human health risks, including cancer, endocrine disruption, and effects on non-target organisms [15–17]. Furthermore, some fungicides have proven ineffective against Foc, necessitating the exploration of alternative, sustainable methods such as biocontrol [18–20].

Biological control involves the use of native microorganisms, known as biocontrol agents (BCAs), to suppress plant pathogens [3, 21]. BCAs are target-specific, environmentally safe, and offer a promising alternative to chemical pesticides [22]. Understanding the modes of action, genetic basis, and biochemical pathways of BCAs is essential for maximizing their efficacy [5]. Despite the identification of secondary metabolites with antifungal properties in several BCAs, their application against Foc remains underexplored, and the regulatory mechanisms of their biosynthesis are poorly understood [23].

In this study, we aimed to isolate and characterize novel beneficial bacterial strains from the rhizosphere of healthy banana plants, for the biocontrol of Foc. To achieve this, we combined multiple complementary approaches. *In-vitro* dual culture assays and GFP-based growth kinetics were used to evaluate the antifungal activity of bacterial isolates against Foc, while greenhouse experiments assessed their protective effects *in-planta*. In parallel, whole-genome sequencing on Illumina platforms [24], followed by comparative genomic and secondary metabolite gene cluster analyses, allowed us to place the most promising isolates within a phylogenetic framework and to predict their capacity to produce antifungal compounds [25, 26]. Joining BCAs from the banana rhizosphere, where they naturally occur, offers a promising approach to mitigate Panama disease within its native ecosystem, reduce disease severity, and increase crop yield.

## 2. Materials and Methods

### 2.1 Isolation of bacteria from healthy banana rhizosphere

Banana root samples were collected from a banana plantation at the Tzemach experimental field, which is situated near the south coast of the Sea of Galilee <u>(32.704545, 35.581159)</u>. No disease symptoms of Foc were observed in the field. For rhizosphere microorganism extraction, an amount of 1 gram of soil samples was mixed into 9 mL of saline solution (0.85%) and mixed twice using FastPrep-24 bead beater (MP Biomedicals, Irvine, CA, USA), at 4.5m/s for 45 seconds and left to settle for 5 min. From the supernatant, 10 µL were spread onto 50% Luria-Bertani (LB50) agar plates consisting of tryptone (5 g/L), yeast extract (2.5 g/L), sodium chloride (5 g/L), and agar (14 g/L) [27] and incubated at 28°C for 2-5 days. For rhizoplane extraction, roots were cut into small fragments, washed with sterile distilled water, and extracted as described above. The resulting suspensions were plated on LB50 agar and incubated under the same conditions. Colonies were separated according to morphological characteristics (colony color, structure, and size). Each colony was purified on a new LB50 plate, followed by growth shaking (150 rpm) in LB50 broth overnight (30°C) and stored in 30% (v/v) of glycerol at -80 °C until used. A total of 436 individual isolates were collected using these methods (see results).

### 2.2 Fungal strains and construction of GFP (green fluorescent protein) labeled Foc

Foc strain was isolated from a symptomatic Cavendish Grande Naine banana plant showing signs of wilting and vascular discoloration. The isolate was tested for pathogenicity in banana plants and remained pathogenic [28]. *Fusarium oxysporum* f. sp. *cannabis* (FC) and *Rhizoctonia solani* (RH) were obtained from the culture collection of Dr. Stanley Freeman, at the Plant Pathology Department, Volcani Center, Agricultural Research Organization (ARO), Israel. Fungal strains were maintained and grown on potato dextrose agar (PDA; Becton, Dickinson and Company, Sparks, USA) medium at 25 °C. For long conservation, the pathogens were kept in 25% (v/v) of glycerol at -80 °C.

Plasmid pCT74 was kindly provided by L. M. Ciuffetti (Oregon State University, Corvallis, Oregon, USA). The vector contains the sGFP gene under control of the ToxA promoter and nos terminator [29]. The Foc TR4-1 isolate was transformed with the GFP-expressing vector using the protoplast method, as previously described [30]. Hygromycin B resistant colonies expressing GFP were collected after 4 days. Single spore isolation was performed and the transformants were checked for stability by transferring on and off PDA medium or PDA Hygromycin B medium (PDA containing 100 μg/μl Hygromycin B) five times. The Foc GFP TR-4 (Foc GFP) isolate was tested twice for pathogenicity in banana plants and remained pathogenic. To further investigate and confirm whether Foc GFP successfully acquired the GFP plasmid, we used confocal microscopy to verify the presence of GFP-labeled *Fusarium* in the banana stem three weeks after inoculating the Foc GFP into the potting mix (Fig. s1).

### 2.3 Evaluation of *in-vitro* antifungal activity by bacterial isolates

The *in-vitro* antagonistic activity of the bacterial strains against Foc, RH, and FC was evaluated using a standard dual culture assay. Twenty microliters of suspension of each of the tested bacterial isolates was streaked in the middle of a 9 cm LB50 agar plates and incubated at 30°C for 1 day. Subsequently, a 4 mm diameter mycelial disk from the margin of an actively growing fungal colony was placed on opposite sides of the Petri dish at a 3-5 mm distance from the edge and further incubated. was conducted at 25°C for 1 week. The area of fungal growth was measured by calculating the diameter of the fungal colony using ImageJ (Wayne Rasband, National Institutes of Health (NIH), Bethesda, Maryland, USA; https://imagej.nih.gov/ij/). The percent inhibition (PI) was calculated using a formula given by Vincent (1947): PI = C-T/C*100, where PI = percent inhibition, C = Radius of Foc colony in the absence of antagonist (cm), and T = Radius of Foc colony in the presence of antagonist bacteria (cm).

### 2.4 *In-planta* greenhouse experiment

Banana plantlets (*Musa* spp., Cavendish subgroup, AAA genome group), propagated via tissue culture, were obtained from Zemach laboratory (Degania, Israel). The plantlets were delivered in plastic seedling pots (3 x 3 x 3 cm) with standard planting medium for banana cultivation. Plantlets ranged in height from 5 to 10 cm and had 4 to 6 leaves.

Bacterial isolates were retrieved from stock cultures stored at -80°C, plated on LB50 agar and incubated overnight at 30°C. The following day, three colonies were selected and suspended in 1000 µl of sterile saline solution. A 100 µl aliquot of this bacterial suspension was spread on three different plates and incubated overnight at 30°C. Bacterial biomass was scraped from LB50 agar plates, resuspended in 2 mL saline, vortexed, and centrifuged (13,000 g, 10 min, 4 °C). The resulting cell pellets were weighed and resuspended in sterile saline (2 mL per 0.1 g of biomass). The suspensions were then adjusted to a final 1:30 dilution relative to the initial biomass suspension and used for inoculation assays. One milliliter of the bacterial suspension was daily applied to the base of the banana plantlets (still in their original seedling cones) for three consecutive days at room temperature, while control plants received 1 ml of water instead.

Forty grams of millet (Maya Foods, Israel) were soaked overnight in sterile water in a 250 ml Erlenmeyer flask. After soaking, the remaining water was discarded, the millet was autoclaved, left at room temperature for 24h, and autoclaved again. A four mm core was taken from PDA plate of 14 days Foc hyphae culture, was then added to the sterile millet and incubated statically for 10-14 days at room temperature without agitation. This culture was incorporated (0.03% w/w) into potting mix and placed in 6.5 cm x 6.5 cm x 8 cm square black pots. Negative control pots contained potting mix with sterile millet.

Before re-planting, plants treated with placed received the third bacterial treatment, and control plants received 1 ml of water or bacterial suspension with no Foc culture. Then, one plantlet with the inoculated cone was planted in each black pot. Pots were placed in plastic containers (6 pots per container) and irrigated by adding water to the bottom of the container until the potting mix was saturated, then transferred to a greenhouse for 40 to 45 days at approximately 28°C and watered daily.

For disease severity index (DSI), the wilt development on each banana plant was determined according to severity scale, as per the following grades: 0 = no disease symptom: the stem is green and healthy, and the leaves are free from discoloration; 1 = slight infection, a minor signs of yellowing appears at the base of the stem, while the rest of the plant, including the leaves, remains healthy;2 = significant yellowing and browning are observed on the stem, and the newly emerging leaf shows yellow streaks; 3 = the entire plant begins to turn yellow, indicating more widespread disease progression; 4 = the plant is completely yellow, and the stem starts bending under its own weight, due to the disease’s impact; and 5 = the plant is entirely brown, dried up, and dead; [32] (Fig. S2).

Disease severity was evaluated at 2-3 days intervals using a 0–5 scale as described above. The area under the disease progress curve (AUDPC) was calculated according to the trapezoidal method [33]:

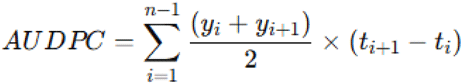

where yi is the disease severity index at the i^th^ observation and ti is the time (days) of the i^th^ observation. Disease suppression by bacterial treatments was expressed as

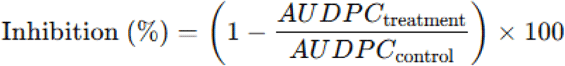

percentage inhibition relative to the pathogen control, using the formula:

where AUDPC_control_ corresponds to plants inoculated with *F. oxysporum* alone, and AUDPC_treatment_ to plants inoculated with both *F. oxysporum* and bacterial strains.

### 2.5 *In-vitro* GFP antifungal activity assay cell-free culture media (CFCM)

For preparing bacterial suspension and CFCM of bacteria, bacteria were inoculated into 4 mL of sterile LB50 in a tube and were cultivated at 27°C, shaking at 170 rpm for 24 h. The cells were pelleted by centrifugation (2400 G, 25°C, 10 min), and the supernatant was filtered through a 0.22 μm pore size filter to remove residual cells or cell debris (Merck Millipore, Darmstadt, Germany) to obtain the CFCM.

To facilitate the production of Foc GFP spores, a nitrate-based sporulation medium was prepared. The medium contained 8.08 g potassium nitrate (KNO_3_), 24 g sucrose, 1.36 g yeast nitrogen base, and 800 mL distilled water. The solution was autoclaved prior to use.

A mycelium-covered agar plug of Foc GFP was transferred into the sterilized KNO_3_ medium and incubated for 48 h. The antifungal activities of CFCM against Foc, were tested using 96-well black, glass bottom plates cell-culture microplates (Greiner Bio-One, Frickenhausen, Germany) in a 96-well microplate reader, Synergy H1 (BioTek, Winooski, VT, USA), as described previously in [34]. Each well of the plate contained 100 μl of potato dextrose broth (PDB; Becton, Dickinson and Company, Sparks, USA) containing 7[×[10^4^ conidia mL^−1^ mixed with CFCM in each well or with 100 μL of LB alone as a positive control. The negative control consisted of 100 μL Foc GFP conidia mixed with 1 μL of 20,000 ppm solution of known translation inhibitor of *Fusarium* growth (Cycloheximide) in 99 μL of LB. All readings were compared to PDB background wells and AUDPC was calculated from the fluorescence data, representing the cumulative fungal growth (as indicated by GFP fluorescence) over time. Lower AUDPC values reflected a stronger inhibitory effect of the bacterial CFCM on Foc GFP growth. Data from at least three independent experiments were used to calculate the average fluorescence.

### 2.6 Microscope and confocal observation with Syto 9 staining

Foc Hyphal morphology was examined using light and fluorescence microscopy. For bright-field observations, samples were collected from the edge of fungal colonies grown either in the absence of bacteria (control) or in dual culture with bacterial treatments. Hyphal morphology was examined using an Olympus light microscope (Olympus CX43, Tokyo, Japan) equipped with an integrated 5-MP digital camera (Olympus EP50) and cellSens imaging software (Olympus, Japan). Observations were performed under bright-field illumination at 40× objectives. Representative micrographs were selected to document hyphal abnormalities, such as swelling, branching, and deformation, induced by bacterial activity.

Fluorescence-based viability imaging of Foc was performed using an Olympus fluoview laser-scanning confocal microscope (Olympus, Tokyo, Japan). The system was equipped with a U-RFL-T fluorescence illumination module, an OBIS multi-line laser unit (Coherent, USA; excitation at 488 nm, 561 nm, and 640 nm), and a motorized objective turret. Imaging was conducted primarily with the 40× water-immersion objective and the 60× oil-immersion objective, depending on the required resolution. Laser power, PMT gain, and offset were adjusted using the Olympus FLUOVIEW acquisition software, maintaining identical acquisition settings for all samples to ensure comparability.

Fungal viability was assessed using the LIVE/DEAD® BacLight™ Bacterial Viability Kit (SYTO 9 / propidium iodide) (Thermo Fisher Scientific, USA). Hyphae or spores were incubated for 30 min in the dark at 25°C with a 1:1 mixture of SYTO 9 (3.34 mM) and propidium iodide (20 mM). SYTO 9 was detected using the 488-nm laser with emission collected at 500–550 nm, whereas propidium iodide was excited using the 561-nm laser with emission collected at 580–650 nm. Z-stacks, XY scans, and transmitted-light channels were recorded using standard confocal modes (XY, XYZ, XYT) of the FLUOVIEW software.

### 2.7 DNA extraction from selected bacterial isolates and genome sequencing

For bacterial identification, the strains were incubated over night at 28°C in LB broth, and fresh cells were harvested by centrifugation. DNA purification was carried out using a Exgene cell SV mini kit (Geneall, Seoul, Korea) following the manufacturer’s instructions according to the manufacturer guidelines. DNA concentration and purity were measured using Qubit (Qubit 2.0 fluorometer, Waltham, MA, USA), and samples were kept at -20°C until further analyzed. Purified DNA was amplified using 16S rRNA universal primers (11 F: 5′-GTTTGATCMTGGCTCAG-3 and 1392 R: 5′-AGCGGCGGTGTGTAC-3′). The reaction was carried out with 5 U Taq DNA polymerase (Thermo Scientific, Waltham, MA, USA) with its reaction buffer according to the manufacturer guidelines. The amplification program consisted of 32 cycles at 95 °C for 30 s, 48 °C for 30 s, and 72 °C for 1 min, and a final extension of 5 min at 72 °C. The amplified product was purified with a PCR purification kit (Thermo Scientific, Waltham, MA, USA) and sequenced (Macrogen, Jerusalem, Israel). Sequences were compared using NCBI BLASTN (http://www.ncbi.nlm.nih.gov) and 98% similarity were considered as identification threshold of these bacterial strains.

Full genome sequencing was performed to the two best candidates (DDC20 and DDC_NEW2). Shotgun sequencing libraries were prepared from bacterial DNA with Illumina DNA Prep (Illumina, 20060059) and sequenced on a NovaSeq X at the Rush University (Chicago, IL, USA).

### 2.8 LC-MS of the bacteria culture medium

Liquid chromatography–mass spectrometry (LC-MS) analysis was used to characterize the compounds potentially involved in the antifungal activity of the culture spent media of strains DDC20 and DDC_NEW2. Aliquots of LB medium (blank control) and culture spent media of strains DDC20 and DDC_NEW2 were filtered through a 0.2 µm PTFE syringe filter, and 5 µL of each sample were injected into an Acquity UPLC system coupled to a Xevo G2-XS QTOF mass spectrometer (Waters Corporation, Milford, MA, USA). Chromatographic separation was performed using a BEH C18 column (2.1 × 50 mm, 1.7 µm) at a flow rate of 0.3 mL min[¹. A linear gradient from water containing 0.5% formic acid (v/v) to acetonitrile containing 0.5% formic acid (v/v) was applied over 30 min. Mass spectra were acquired in both positive and negative electrospray ionization (ESI) modes, with the positive mode providing the strongest ion signals for metabolites present in the culture media (Fig. S4 & S5). By visualizing the final chromatographic profiles of each strain, major classes of metabolites were detected, and compared.

### 2.9 Extraction of metabolites and preparation of crude extracts for antifungal assays

To evaluate whether metabolites produced by the bacterial strains were influenced by the presence of live or dead (heat-killed) fungal biomass, an elicitation assay was performed. The DDC20 and DDC_NEW2 strains were grown under three culture conditions: (a) no fungal addition, (b) live fungi addition (LF), (c) dead fungi addition (DF), and the produced metabolites were extracted as follows. Fifty mL of each spent culture medium was first extracted with ethyl-acetate (EA) (3 x 25 mL) and then with n-BuOH (BuOH) (30 mL). The extracts were dried and weighed. The resulting extracts were analyzed by LC-MS as described above (2.8) and used for antifungal analysis. All dried extracts (EA and BuOH) were stored at -20°C and resuspended in DMSO prior to antifungal bioassays.

For antifungal assays, dried EA and BuOH were resuspended in dimethyl sulfoxide (DMSO) to a final concentration of 20 mg mL[¹. The extracts were vortexed thoroughly to ensure complete dissolution, sterile-filtered through a 0.22 µm PTFE syringe filter and stored at –20°C until use. Immediately before testing their antifungal activities, working dilutions of the crude extracts were prepared in PDB to match the final DMSO concentration (5%) across all treatments, including control wells.

### 2.10 GFP-based antifungal assay using crude organic extracts

The antifungal activity of the crude extracts was evaluated using the same GFP-based microplate assay described for the CFCM (Section 2.5). In each well, 100 µL of PDB containing 7 × 10[ Foc GFP conidia mL[¹ were mixed with the appropriate dilution of the extracts to reach final test concentrations equivalent to those used for CFCM. Fluorescence kinetics were monitored every 30 min for 92 h at excitation/emission wavelengths of 479/520 nm using a Synergy H1 microplate reader. AUDPC values were calculated to quantify cumulative fungal growth inhibition. Crude-extract treatments were directly compared to CFCM, DMSO control (5%), cycloheximide control (10 ppm), and positive-growth controls (no treatment).

### 2.11 Data analysis

All experiments were conducted in three biological replicates, and data were presented as the mean ± standard deviation. Initial *in-vitro* experiments visually assessed the inhibitory qualitative effects of bacterial strains against Foc. For quantitative analysis, an analysis of variance (ANOVA) model was applied to each experimental setup to evaluate the significance of the effects. Post-hoc Dunnett’s tests (α=0.05) were used to compare the inhibitory effects of individual bacterial strains with that of the control group.

Greenhouse experiments were conducted using a randomized design. An ANOVA model was employed to analyze the effects of bacterial strains on disease progression, with post-hoc Dunnett’s tests (α=0.05), identifying strains that significantly reduced AUDPC values compared to the control group.

To evaluate the inhibitory effects of CFCM from bacterial strains against Foc GFP spores, fluorescence-based AUDPC data were log-transformed to stabilize variances and normalize the data distribution. This transformation minimized the impact of outliers and ensured the assumptions of the ANOVA model were met. An ANOVA model was then applied to compare the effects of bacterial CFCM treatments on Foc GFP growth. Post-hoc Dunnett’s tests (α=0.05) were used to identify bacterial strains with supernatants that significantly reduced AUDPC values relative to the control.

For the two selected top candidates, an additional ANOVA model was used, treating bacteria as a fixed effect, while the experimental setup and the interaction between bacteria and experimental conditions were treated as random effects. The Wald p-value for the interaction between the experimental setup and treatment was examined to determine whether the bacterial effects remained consistent across experimental conditions. Contrast t-tests were further conducted to compare each bacterium against Foc.

All statistical analyses were performed using JMP pro (18.0.2) software (SAS Institute, Cary, NC).

## 3. Results

### 3.1 Screening and identification of bacteria with antifungal activity

A total of 436 bacterial strains were isolated from the rhizosphere soil of a healthy banana plant. Of these isolates, 123 showed some inhibition activities against Foc on LB50 agar plates, and the activity of 93 of them was statistically significant. To identify the most effective strains, 78 bacterial strains with the strongest anti-Foc *in-vitro* activity were further evaluated. Cell-free culture media (CFCM) from these strains were tested for inhibitory effects against Foc GFP spores. Statistical analysis revealed that 48 CFCM significantly reduced Foc GFP spore germination compared to the control.

To identify the best-performing isolates *in-planta*, strains that showed high *in-vitro* activity were also tested for their ability to reduce disease severity in greenhouse experiments. These experiments were conducted using banana plants (*Musa spp., Cavendish subgroup,* AAA genome group) inoculated with Foc. Disease progression (DSI from 0 (healthy plant) to 5 (dead plant)) was monitored for 45 days (Fig. S2). To quantify disease progression over time, the area under the disease progression curve (AUDPC) was calculated. The AUDPC provides an integrated measure of disease intensity over the experimental period, reflecting the cumulative impact of bacterial treatment on disease suppression. A lower AUDPC value indicates greater efficacy of the bacterial strain in mitigating disease severity compared to the control.

The results revealed that 22 bacterial strains demonstrated a statistically significant reduction in AUDPC, confirming their potential Foc-induced disease inhibition in banana plants under greenhouse conditions.

Ultimately, in the selection process described so far, two isolates (DDC20 and DDC_NEW2) were selected as best candidates due to their consistently high performance in all assays as will be discussed below (Fig. 1, Fig 2). In the *in-vitro* assays, which tested bacterial inhibition zone versus Foc, the mean Foc growth inhibition diameters for strains DDC_NEW2 and DDC20, were 9 and 8.50 mm (respectively), significantly smaller than no-bacteria (22.07 mm) and *E. coli* (21.19 mm) controls. These differences were highly significant (p ≤ 0.0002 in all cases), demonstrating the strong inhibitory activity of these strains (Fig. 1A and B). For the two selected bacterial candidates, data from three independent *in-vitro* Petri dish experiments performed on different dates were analyzed using a mixed-effects ANOVA model. The interaction between bacterial treatment and experimental setup was included to test whether bacterial effects varied across experiments. This interaction was not statistically significant (p > 0.05), indicating that the inhibitory effects of the bacteria were consistent and reproducible across independent experimental runs.

**Fig. 1:**
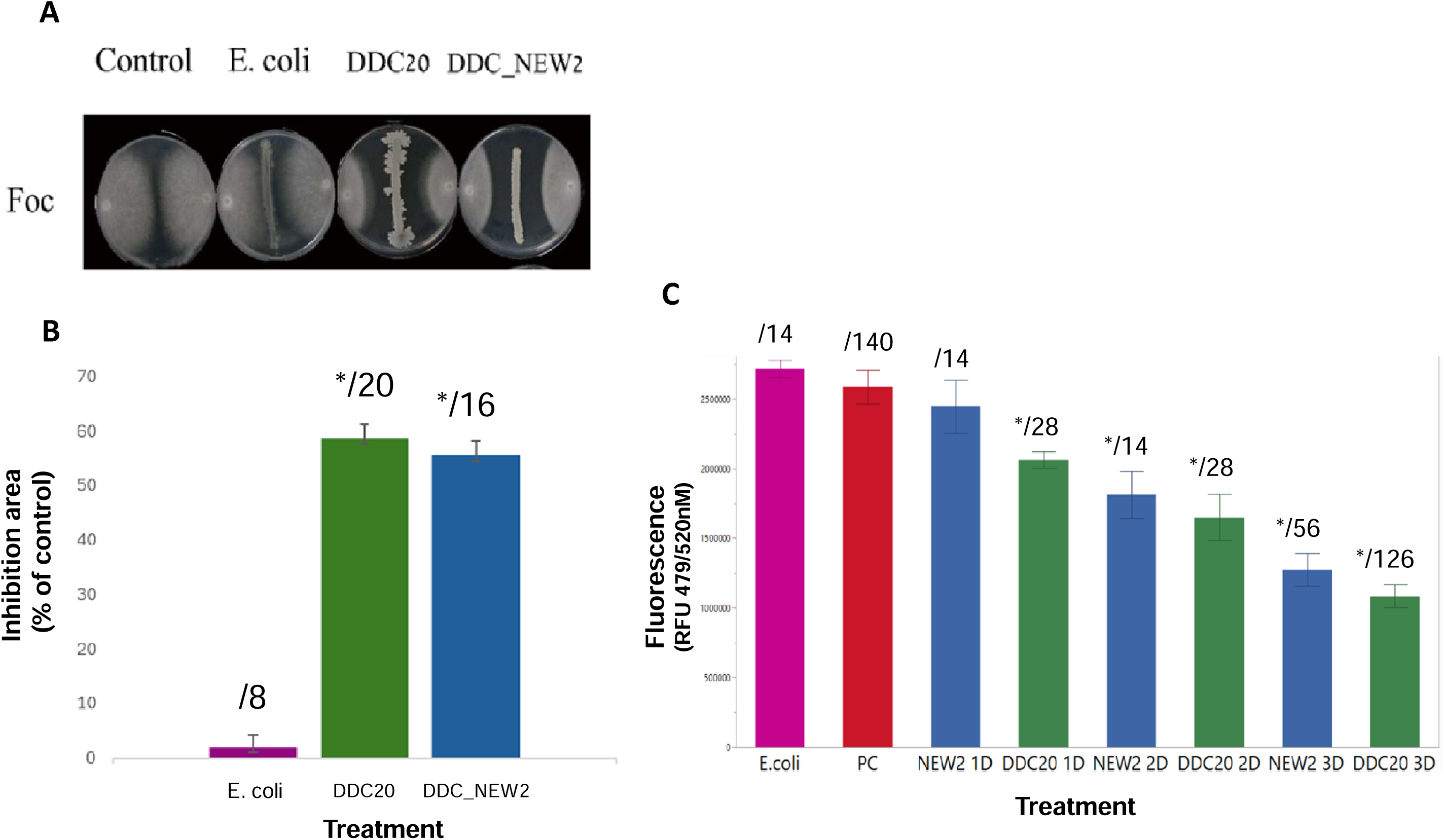
In vitro inhibition of Fusarium oxysporumf. sp. cubense(Foc) by antagonistic bacteria. (A) Agar plate culture assay showing the growth of Foe against no-bacteria control, E. coli (negative control), and the antagonistic strains DDC2O and DDC_NEW2. (B) Percent inhibition area of Foe mycelial growth by the bacterial strains compared to the control (calculated from A). (C) Effect of bacterial culture supernatants on F. oxysporum f. sp. Cubense GFP (Foc-GFP) spore germination. Fluorescence was calculated for the cell-free culture media (CFCM) of bacterial strains E. coli, DDC2_NEW2 and DDC2O, during incubation periods of 1 (ID), 2(2D), 3 (3D) days. The x-axis shows different treatments, while the y-axis represents fluorescence values. The numbers above each bar indicate the sample size (N)for each treatment, error bars indicate standard error of the mean. Asterisks (*) denote statistically significant differences relative to the control according to Dunnett’s test (P < 0.05).

**Fig. 2:**
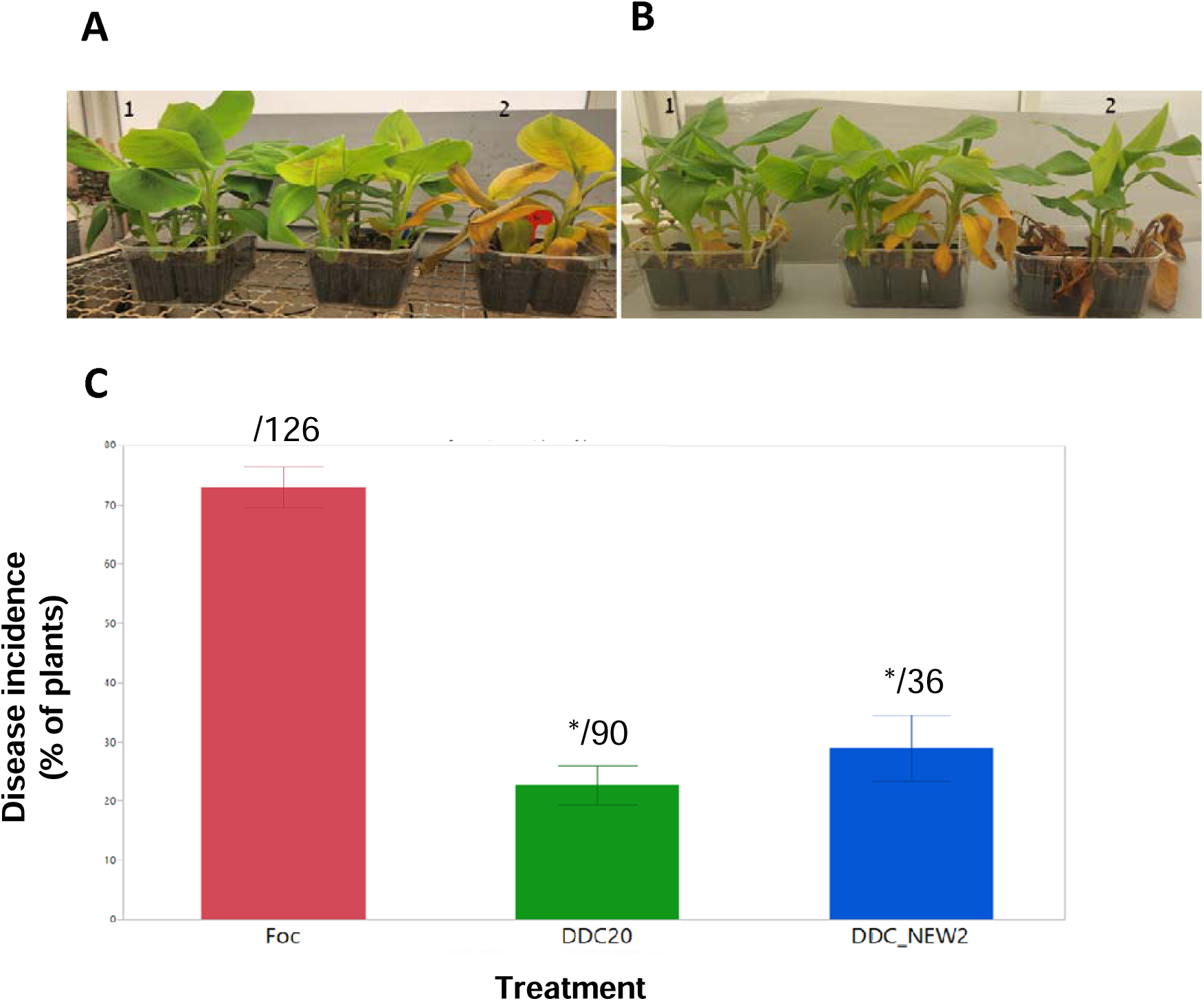
Effect of bacterial treatments on banana plants infected with Fusarium oxysporumf. sp. cubense (Foe). Effect of (A) DDC2O treatment and (B) DDC_NEW2 treatment. Number 1 = negative control (healthy plants grown without pathogen); 2 = Positive control (plants inoculated with Foe with no bacterial addition (C) The graph represents the inhibition percentages of bacterial isolates DDC2O, and DDC_NEW2 against our control Foe in greenhouse, by using disease severity scale. Error bars indicate standard deviation. Statistical significance was determined using Dunnett’s test (p < 0.05), indicated by an asterisk (*). The numbers above each bar indicate the sample size (N) for each treatment.

To further evaluate the robustness and broad-spectrum inhibitory potential of these bacterial strains, we tested their inhibition activity against other fungi: FC and RH. The results demonstrated that DDC_NEW2 and DDC20 exhibited inhibitory effectiveness against FC and to a lesser extent also against RH (Fig. S3).

On the CFCM effectivity assay (Fig. 1C), DDC20’s CFCM consistently exhibited a significantly higher Foc inhibition compared to the control (Foc only) (p ≤ 0.003), with a mean AUDPC of 1.36 × 10[ versus 2.57 × 10[ for the control. In contrast, the differences between DDC_NEW2 (mean AUDPC: 1.54 × 10[) against Foc was inconsistent and not statistically significant (p = 0.17). Analysis of the interaction between the experimental setup and treatment parameters in this step was statistically significant (p ≤ 0.001), suggesting variability in bacterial effects across experiments. The results of DDC_NEW2 and DDC20 in the *in-planta* experiment (Fig. 2), were similar to the *in-vitro* results. The mean AUDPC values for DDC20 and NEW2, were 22.7 and 28.9, respectively, which are significantly (p ≤ 0.0002) smaller than that of Foc (73) (Fig. 2C). Analysis of the overall interaction between the experimental setup and treatment parameters was not statistically significant (p > 0.05), confirming consistent bacterial effects across experimental conditions. These highly significant results indicate that these bacterial strains could effectively reduce disease severity *in-planta* as well as *in-vitro*.

### 3.2 Comparative Microscopic Analysis

Next, to further characterize the mode of action of DDC20 and DDC_NEW2, we examined the morphology of Foc hyphae from the *in-vitro* petri dish assay, after exposure to the bacterial strains *in-vitro* (Fig. 3). Microscopic analysis revealed striking morphological differences between the control, non-treated cultures and those co-cultivated with the bacterial strains. In the absence of bacterial antagonism, Foc produced typical hyphal networks with well-formed macroconidia (3–5 septa) and microconidia, displaying the expected fusiform morphology and uniform growth pattern. By contrast, in the presence of either DDC20 or DDC_NEW2, Foc exhibited pronounced hyphal abnormalities, including swelling, irregular branching, and distorted growth. Conidial development was altered, with irregularly distributed macroconidia and microconidia often associated with malformed hyphae. Notably, exposure to both bacteria triggered abundant chlamydospore formation, a hallmark of stress and fungal survival under unfavorable conditions. At higher magnification, deformed hyphae with intercalary swellings and clusters of thick-walled chlamydospores were evident, confirming that exposure to the bacteria exerts a strong stress effect on fungal development.

**Fig. 3:**
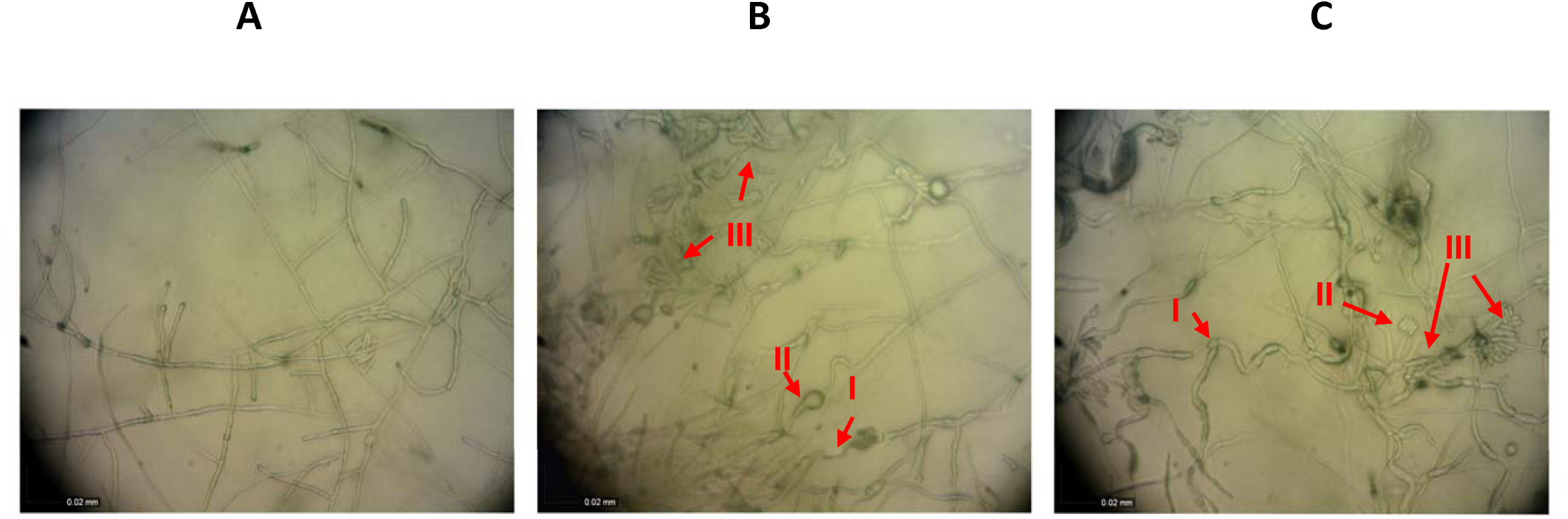
Microscopic observation of Fusarium oxysporum f. sp. cubense (Foe) hyphae in the presence and absence of bacteria. Light microscopy showing Foe hyphal morphology under different treatments: (A) no bacteria control; (B) exposure to DDC2O during seven days; (C) exposure to DDC_NEW2 during seven days. Numbers indicate severe hyphal abnormalities, including distortion (I), intercalary swellings (II), and abundant sporulation with clusters of microconidia, macroconidia, and chlamydospores (III).

Overall, the interaction of Foc with DDC20 or DDC_NEW2 resulted in suppressed mycelial growth, aberrant sporulation, and structural deformations consistent with potent antifungal activity.

### 3.3 Confocal Microscopy Observations of Cell-Free Culture Media (CFCM) effects

To further investigate the biocontrol potential of DDC_NEW2 and DDC20 against Foc, we also performed confocal microscopy observations on hyphae exposed to CFCM supplemented after staining with the fluorescent dyes SYTO 9/ propidium iodide (PI) (Fig. 4). SYTO 9/PI staining can be used to assess cell viability, as SYTO 9 penetrates intact cell membranes and stains live cells in green [35, 36] while, PI can penetrate dead cells with compromised membranes, and stain them in red. Our results revealed a significant difference between the hyphae exposed to DDC_NEW2 and DDC20 treatment and the control (hyphae not exposed to the bacteria). In the presence of both strains CFCM, and especially with DDC20, many hyphae and spores were stained red, indicating cell damage and possible cell death compared to the control, where fungal structures remained largely viable. These findings suggest that the bacterial strains exert a direct or indirect inhibitory effect on Foc, leading to fungal cell stress and cell death, reinforcing their potential as biocontrol agents in banana plants.

**Fig. 4:**
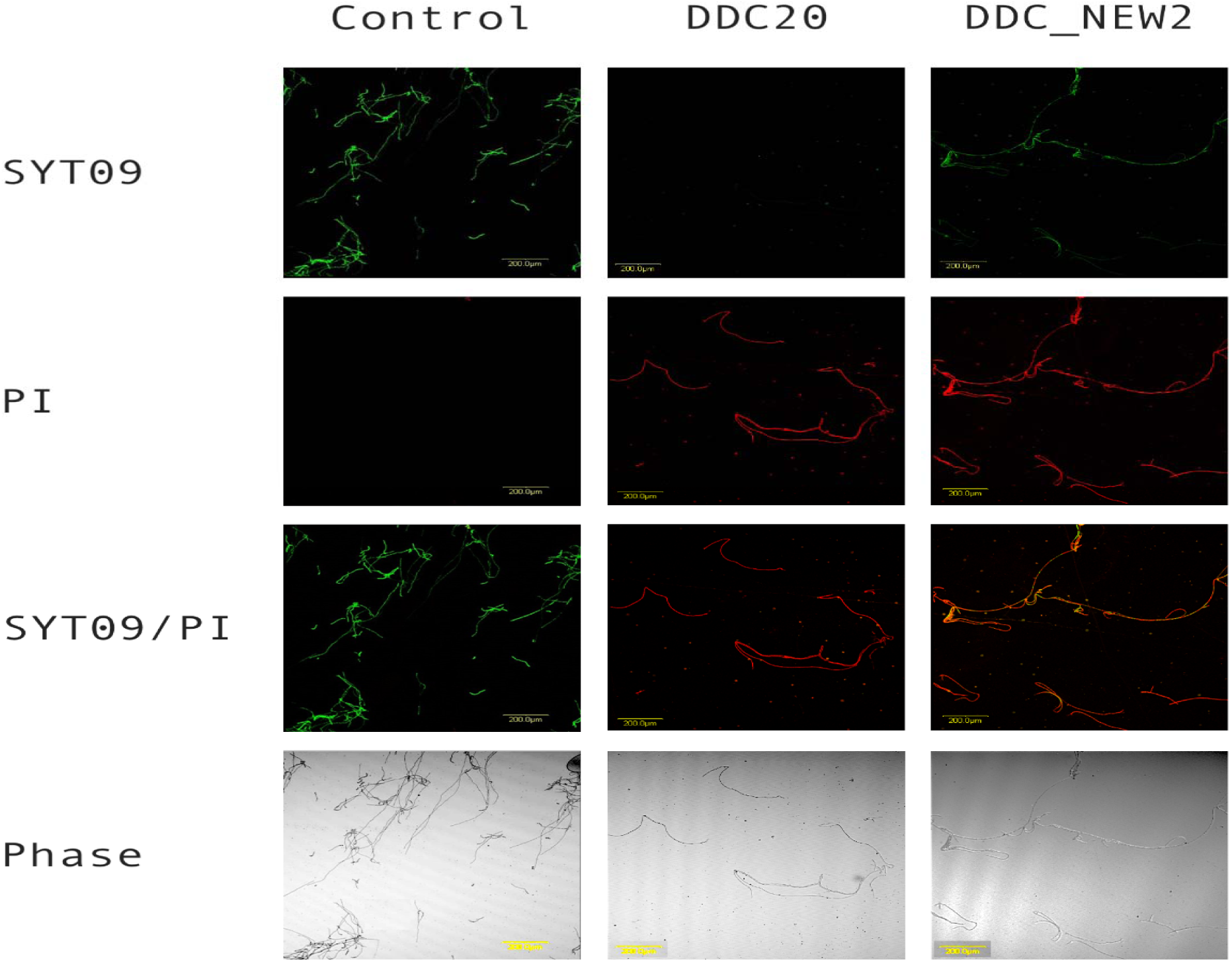
Confocal laser scanning microscopy showing viability loss of Fusarium oxysporum f. sp. cubense (Foe) hyphae after exposure to cell-free culture media (CFCM). Foe hyphae were treated with CFCM from strains DDC2O and DDC_NEW2 and stained with SYTO 9 (green, viable cells) and propidium iodide (PI; red, membrane-compromised cells). Control samples show intact, strongly fluorescent green hyphae, indicating high viability. DDC2O and DDC_NEW2-treated hyphae exhibit a drastic reduction in SYTO 9 signal and intense PI staining, reflecting severe membrane damage and loss of viability. Merged SYTO9/PI images confirm the shift from viable (green) to dead/damaged (red) hyphae following treatment. Phase-contrast images show morphological deformation consistent with membrane disruption. Scale bar = 200 µm.

### 3.4 Morphology and Genomic Features and Phylogenetic Position of *isolates* DDC20 and DDC_NEW2

On agar plates, the colonies of DDC20 exhibited a light yellow, slightly irregular morphology, while colonies of DDC_NEW2 displayed a viscous creamy-white pigmentation with a smooth margin. DDC20 and DDC_new2 were classified as Gram-positive, motile, endospore-forming, and catalase-positive bacteria (data not shown).

The genomes of strains DDC20 and DDC_NEW2 were sequenced, assembled and analyzed using QUAST. The two bacteria have been phylogenetically characterized and positioned within the *Bacillus* clade, closely related to *B. halotolerans*, *B. subtilis, B. amyloliquefaciens* and *B. velezensis* (Fig. 5). These analyses suggest that DDC20 and DDC_NEW2, although phylogenetically distinct, are both related to well-characterized plant-associated *Bacillus* species. Members of these species are known for their roles in the suppression of plant pathogens and in plant growth promotion. Genomic analyses reveal that DDC20 and DDC_NEW2 possess circular chromosomes with genome sizes of 4.12 Mb and 4.32 Mb and GC contents of 43.72 % and 46.39 %, respectively. These values are consistent with those observed in closely related *Bacillus* species, which typically range from 3.38 Mb to 4.96 Mb with GC contents between 43.8% and 46.8%. The genome of DDC20 was assembled into 21 contigs and DDC_NEW2 into a total of 62 contigs. For DDC20 the largest contig is 1,086,518 bp. The assembly is more contiguous, as indicated by a higher N50 value of 1,065,504 bp and an L50 of 2, demonstrating that two large contigs make up 50% of the genome. For DDC_NEW2 the largest contig in this assembly measures 1,068,876 bp, slightly shorter than that of DDC20. The assembly quality is reflected in an N50 value of 994,374 bp and an L50 value of 2, indicating that two large contigs account for at least 50% of the total genome size. A lower percentage of gaps in DDC20 (0.005%) compared to DDC_NEW2 (0.017%) further supports the higher contiguity of DDC20’s genome. To complement the assembly quality evaluation, BUSCO analysis was performed to assess genome fullness using a benchmark set of conserved single copy orthologs (Supplementary Fig. 5). Both DDC_NEW2 and DDC20 exhibit 100% completeness, with all 124 expected BUSCO genes present and intact, confirming the high quality and near-completeness of both genome assemblies. Notably, both genomes contain 124 complete single-copy BUSCO genes and 0 duplicated, fragmented, or missing genes, reinforcing the reliability of these assemblies (Tab. S1).

**Fig. 5:**
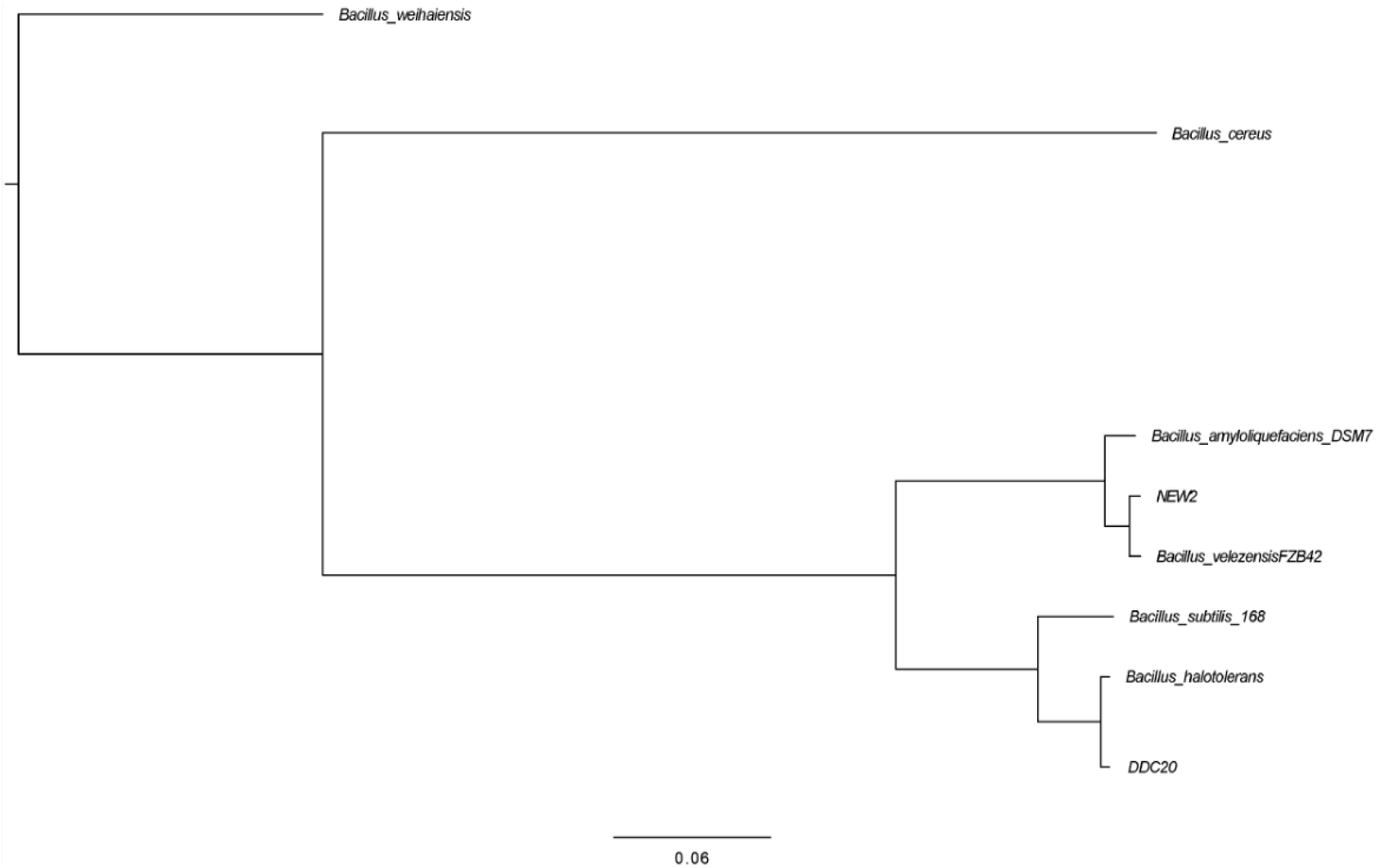
Phylogenetic tree of Bacillus strains, including isolates DDC2O and DDC_NEW2. The tree represents the phylogenetic relationship among selected Bacillus species based on full genomic sequence analysis. The scale bar represents the substitution rate per site.

### 3.5 LC-MS analysis and elicitation assays

Next, LC-MS profiling was conducted to determine whether the presence of *F. oxysporum*, either alive or heat-killed, elicits metabolite production in strains DDC20 and DDC_NEW2. Across all LC-MS chromatograms, three retention-time regions were reproducibly observed: an early low-intensity segment between 1.5 and 8.0 min corresponding to the LB medium components; a mid-retention zone (12–15 min) dominated by fengycin homologs ions (*m/z* 1449–1505); and a late region (20–23.5 min) containing surfactin homologs ions (*m/z* 1008–1036) (Fig. S4, Fig. S5). No additional peaks were detected when cultures were supplemented with either live or dead Foc, demonstrating that neither condition induced nor repressed lipopeptide biosynthesis.

Although both strains shared the same qualitative families of compounds, their quantitative and compositional profiles markedly differed. DDC_NEW2 displayed a strong and coherent fengycin cluster with major ions at Rt 13.12 min (*m/z* 1449.7880), Rt 13.85 min (*m/z* 1463.8043), Rt 14.30 min (*m/z* 1477.8024), and Rt 15.15 min (*m/z* 1505.8505) (Fig. 7). In contrast, DDC20 produced fengycins at a lower concentration but exhibited a broader metabolite repertoire, including a distinct mojavensin signal at Rt 12.73 min (*m/z* 1106.5614) and a pronounced group of unidentified metabolites eluting between 9.5 and 11 min (features that were nearly absent from the DDC_NEW2 chromatogram) (Fig. 6). The surfactins region was similar in both strains, with homologues eluting at Rt 21–23.5 min, although relative peak intensities varied moderately between strains.

**Fig. 6:**
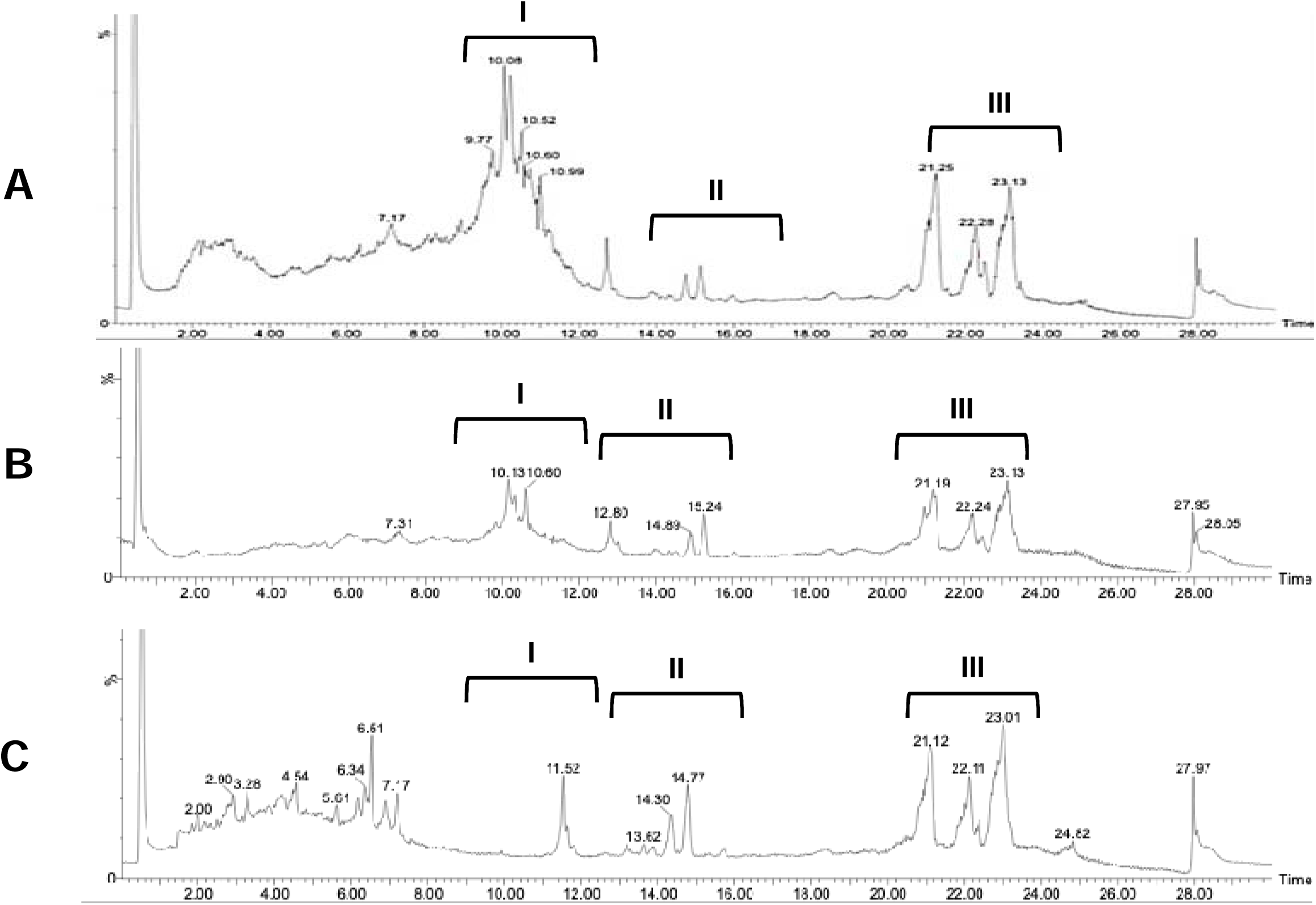
LC-MS total ion chromatograms (ESI*) of DDC2O culture supernatants grown without Foe (A), or in the presence of live (B) or heat-killed Foe (C). Chromatographic profiles consistently show three major retention regions: a first region with several unidentified metabolites eluting at 9.5-11 minutes (I); a second region between 12 and 15 minutes corresponding to thefengyein family (II); and a third region between 21 and 23.5 minutes containing surfactin homologues (III). A mojavensin-associatedpeak (Rt ∼I4.7 min) is detected in all chromatograms.

**Fig. 7:**
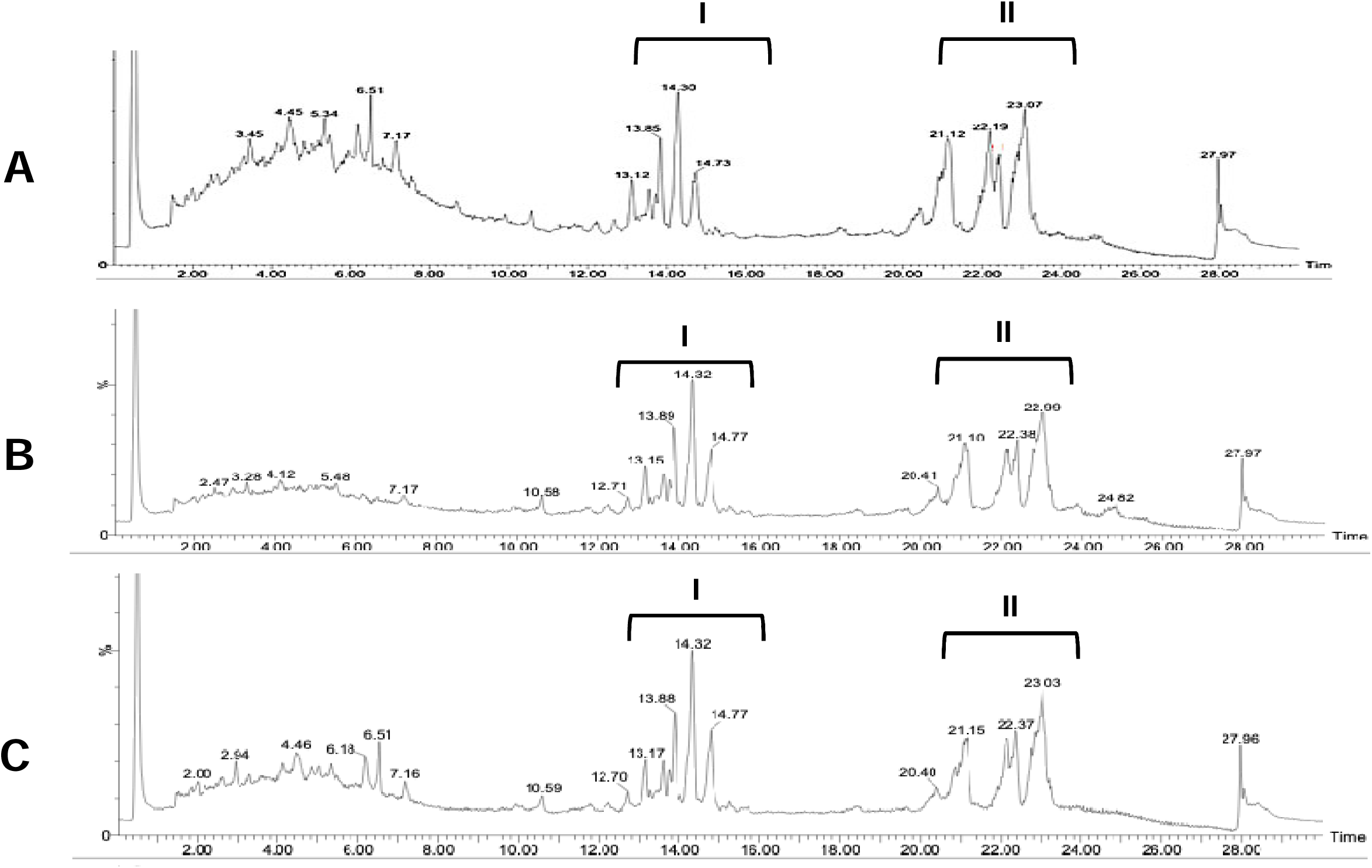
LC-MS total ion chromatograms (ESI*) of culture supernatants of DDC_NEW2 grown without Foe, or in the presence of live or heat-killed Foe. All chromatograms, (A) without Fusarium oxysporum (no Foe), (B) with live Foe, and (C) with heat-killed Foe display the same characteristic retention regions. Two main groups are detected: dominant fengyein cluster between 12 and 15 min (I) and surfactin homologues eluting between 21 and 23.5 min (II).

Altogether, LC-MS analyses demonstrate that DDC20 and DDC_NEW2 constitutively produce fengycins and surfactins, and DDC20 exhibited a wider and more complex metabolic profile, including mojavensin and several unidentified metabolites. The stability of these profiles across elicitation conditions supports the conclusion that antifungal lipopeptide production is intrinsic to both strains rather than triggered by fungal presence.

To determine which classes of compounds, contribute to the antifungal activity of strains DDC20 and DDC_NEW2, bacterial cultures were extracted using EA or BuOH. Since EA extracts were shown (by LC-MS) to contain only the surfactins, while the BuOH extracts were shown to contain the more polar fengycins and LB-medium components, both were tested for their ability to inhibit GFP-expressing Foc. Eight samples were evaluated: four derived from DDC_NEW2 (A: EA extract of dead fungi; B: EA of extract of live fungi; C: BuOH-extract of dead fungi; D: BuOH of live fungi) and four from DDC20 (E, EA extract of dead fungi; F, EA of extract of live fungi; G, BuOH-extract of dead fungi; H, BuOH of live fungi), Across all treatments, inhibition levels were quantified by measuring fluorescence kinetics and calculating the AUDPC. Fractions A–D, originating from DDC_NEW2, displayed the lower antifungal activity then DDC20. DDC_NEW2 EA extracts, (A and B) produced only slight reductions in AUDPC compared with the positive control. BuOH extracts (C and D), which are enriched in more polar compounds such as fengycins, resulted in stronger inhibition than EA fractions, consistent with LC–MS profiles. Interestingly, combinations of DDC_NEW2 fractions revealed additive effects. For example, fractions A + C inhibited fluorescence more strongly than either A or C alone, and B + D exceeded the inhibition of B and D individually. These patterns suggest that apolar and polar metabolites in DDC_NEW2 act additively when combined (Fig. 8). Fractions E–H from DDC20 extract showed a stronger inhibition profile. EA fractions (E and F) displayed higher antifungal activity than any DDC_NEW2-derived EA extract, consistent with the higher concentration of surfactins measured DDC20 extract. BuOH fractions, (G and H), produced the strongest suppression of GFP fluorescence among all tested samples, with AUDPC values approaching complete inhibition (Fig. 8). Due to the very high inhibitory activity of each of the DDC20 fractions, combining fractions did not reveal additional suppressive synergistic effects (reaching maximal “ceiling” inhibition). This strong inhibitory performance of fractions G and H aligns with LC–MS measurements showing abundant polar lipopeptides (fengycins) and several unidentified metabolites present uniquely in this fraction of DDC20 extract. No crude-extract treatment exceeded the inhibition observed with the cycloheximide control.

**Fig. 8:**
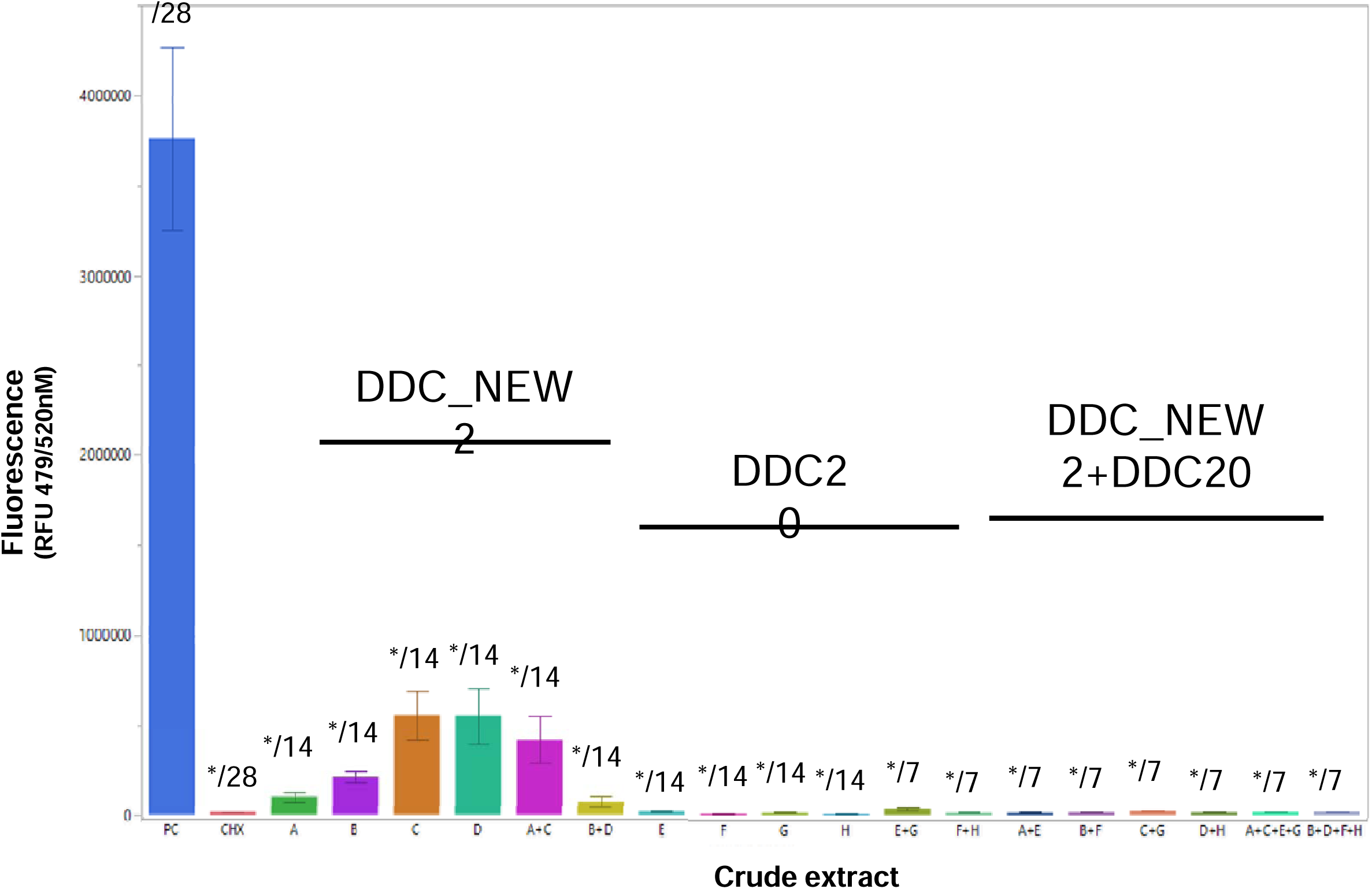
Antifungal activity of chemically fractionated extracts from strains DDC_NEW2 and DDC2O against GFP-Foc. Fluorescence-based inhibition of GFP-Foc after treatment with individual or combined chemical extracts derived from DDC_NEW2 (fractions A-D), DDC2O (fractions E-H, same), or mixtures of specific extracts from both strains. Extracts were obtained following solvent partitioning (ethyl acetate, EA; n-butanol, BuOH) of culture supernatants prepared from cultures exposed to either live (LF) or heat-killed (DF) fungal biomass. DDC_NEW2-derivedfractions (A-D) showed modest inhibition, whereas DDC2O fractions (E-H) displayed strong to nearly complete suppression of fluorescence. All combinations containing at least one DDC2O fraction consistently retained high inhibitory activity. PC: positive control (Foe GFP without treatment). CHX: cycloheximide (5 ppm, negative inhibition control). The numbers above each bar indicate the sample size (N) for each treatment, error bars indicate the standard error of the mean. Asterisks (*) denote statistically significant differences relative to the control according to Dunnett’s test (P < 0.05).

Overall, these results demonstrate that CFCM produced a measurable but moderate reduction in AUDPC compared to the positive control (Fig. 1). Crude organic extracts exhibited substantially stronger inhibitory effects, with several treatments reducing AUDPC by more than 70–90%. DDC20 extracts consistently outperformed those of DDC_NEW2 by reducing AUDPC by more than 99% compared to the control (Fig. 8). DDC20 produces a more potent and compositionally richer combination of antifungal metabolites, which is consistent with its superior performance in-vitro, in-planta, and in LC–MS chromatographic analyses.

### 3.6 Functional Classification and Its Relation to Biosynthetic Potential

Genomic functional classification of DDC_NEW2 and DDC20 revealed that Clusters of Orthologous Groups (COG) had broadly similar distribution patterns (Fig. 9). Both genomes were enriched in categories related to primary metabolism and cellular processes, with the most represented being E (amino acid transport and metabolism, ∼9%), G (carbohydrate transport and metabolism, ∼8–9%), and K (transcription, ∼8%). Other categories with substantial representation included J (translation, ribosomal structure and biogenesis, ∼7%), D (cell cycle control, cell division, chromosome partitioning, ∼7%), and in DDC20 also M (cell wall/membrane/envelope biogenesis, ∼6%).

**Fig. 9:**
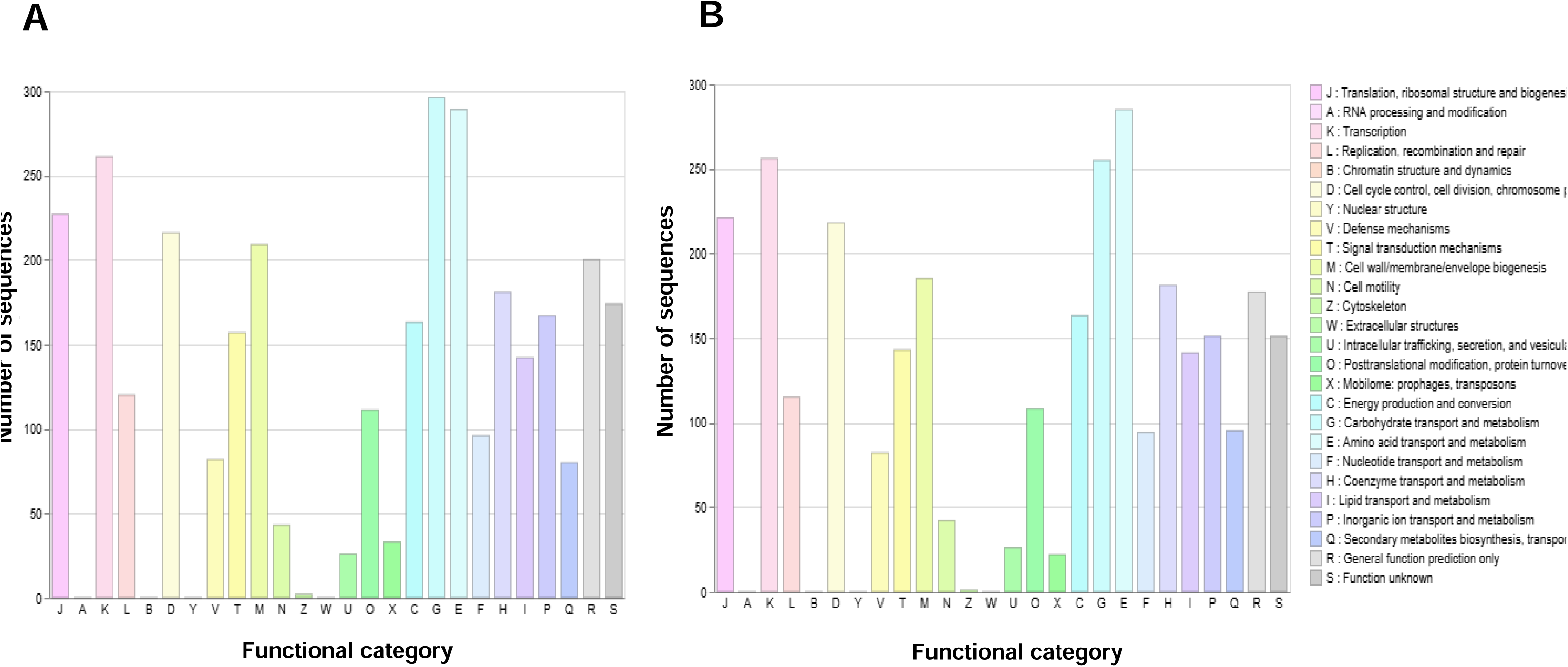
COG functional classification of predicted protein-coding genes. Functional category for DDC2O (A) and DDC_new2 (B). Each bar represents a functional category (A to S), with the number of sequences indicated on the Y-axis. The clusters of orthologous groups (COG) categories highlight the distribution of gene functions across major biological processes, including secondary metabolite biosynthesis (Q).

In contrast, categories such as N (cell motility), U (intracellular trafficking, secretion, and vesicular transport), X (mobilome, prophage, transposons), and Z (cytoskeleton) were poorly represented in both strains, each with fewer than 50 sequences. Notably, a considerable proportion of genes in both genomes were classified as R (general function prediction, ∼6–7.5%) and S (function unknown, ∼5–6%), indicating that a substantial portion of their coding potential remains uncharacterized.

Overall, the COG profiles of DDC_NEW2 and DDC20 emphasize their metabolic versatility while also highlighting the presence of numerous genes of unknown function that could contribute to their biocontrol activity.

The presence of secondary metabolite-related genes (Q category) in both strains was verified using antiSMASH analysis (Antibiotics and Secondary Metabolite Analysis Shell) tool (v7.1.0) (Table 1&2). antiSMASH identifies multiple biosynthetic gene clusters (BGCs) involved in antibiotic and antifungal compound biosynthesis.

**Table 1:** Secondary metabolite biosynthetic gene clusters predicted in the genome of the bacterial isolate using antiSMASH. . The table lists the contig, region, cluster type, genomic coordinates, the most similar known cluster, compound class, and percentage similarity. This table represents secondary metabolites producing biosynthetic clusters of DDC_NEW2 with homology to the most similar known biosynthetic clusters as predicted by antiSMASH.

| Contig | Region | Type | Start | End | Most Similar Known Cluster | Cluster Type | Similarity (%) |
| --- | --- | --- | --- | --- | --- | --- | --- |
| contig_1 | Region 1.1 | NRPS, bacteriocin, NRPS-like, RiPP-like | 73719 | 132362 | bacillibactin | NRP | 100 |
| contig_1 | Region 1.2 | other | 651416 | 692837 | bacilysin | Other | 100 |
| contig_2 | Region 2.1 | NRPS | 1 | 14454 | fengycin | NRP | 13 |
| contig_4 | Region 4.1 | PKS-like | 63428 | 104672 | butirosin A/butirosin B | Saccharide | 7 |
| contig_4 | Region 4.2 | terpene | 186685 | 207425 |  |  |  |
| contig_4 | Region 4.3 | transAT-PKS | 502181 | 590396 | macrolactin | Polyketide | 100 |
| contig_4 | Region 4.4 | transAT-PKS, NRPS, T3PKS | 809198 | 919315 | bacillaene | Polyketide+NRP | 100 |
| contig_4 | Region 4.5 | NRPS, transAT-PKS, beta-lactone | 980876 | 1068876 | fengycin | NRP | 80 |
| contig_9 | Region 9.1 | terpene | 45879 | 67762 | - | - | - |
| contig_9 | Region 9.2 | T3PKS | 136424 | 177524 |  |  |  |
| contig_9 | Region 9.3 | transAT-PKS-like | 292949 | 339537 | difficidin | Polyketide | 53 |
| contig_12 | Region 12.1 | NRPS | 1 | 25205 | surfactin | NRP-Lipopeptide | 39 |
| contig_13 | Region 13.1 | NRPS, transAT-PKS | 1 | 46600 |  |  |  |
| contig_13 | Region 13.2 | NRPS | 115894 | 144172 | surfactin | NRP-Lipopeptide | 47 |
| contig_14 | Region 14.1 | lanthipeptide-class-iii | 21372 | 43987 | andalusicin A/andalusicin B | RiPP:Lanthipeptide | 100 |
| contig_15 | Region 15.1 | transAT-PKS-like | 1 | 23789 | difficidin |  | 26 |
| contig_17 | Region 17.1 | NRPS | 1 | 9574 | surfactin | NRP-Lipopeptide | 8 |
| contig_18 | Region 18.1 | NRPS | 1 | 6522 |  |  |  |
| contig_20 | Region 20.1 | NRPS | 1 | 10019 |  |  |  |

The analysis revealed diverse groups of BGCs in both bacterial genomes, including nonribosomal peptides (NRPS), polyketides (PKS), ribosomally synthesized and post-translationally modified peptides (RiPPs), terpenes, and β-lactones.

DDC_NEW2 has a total of 19 BGCs detected across multiple contigs (Table 1). Several NRPS-related clusters were identified, with the most prominent being associated with the production of bacillibactin (100% similarity), bacilysin (100%), fengycin homologues (80% and 13%), and surfactin homologues (47,39 and 8%). These lipopeptides and siderophores play a crucial role in iron scavenging, antimicrobial activity, and plant protection, suggesting that the DDC_NEW2 can produce potent antibacterial compounds, and be a potential biocontrol agent against fungal pathogens such as Foc.

Several polyketide synthases (PKS) and transAT-PKS clusters were detected, including a difficidin-like cluster (53% similarity), which is known for its antibacterial properties. The transAT-PKS-like clusters, found in contig_9 and contig_15, suggest that DDC_NEW2 might produce additional metabolite derivatives with antimicrobial activity. A lanthipeptide-class-III cluster (100% similarity to andalusicin A/B) was identified in contig_14, suggesting DDC_NEW2 can produce antimicrobial RiPPs with potential pharmaceutical applications. Lanthipeptides are known for their broad-spectrum antibacterial activity, making this cluster of particular interest. Additionally, terpene biosynthesis genes were found in contig_4 and contig_9, though their functional products remain uncharacterized. The presence of a beta-lactone biosynthesis gene cluster suggests potential protease inhibition or antimicrobial activity.

Genome analysis of strain DDC20 using antiSMASH revealed also a diverse repertoire of biosynthetic gene clusters (BGCs), indicating a substantial potential for secondary metabolite production (Table 2). Several predicted clusters showed high similarity to well-characterized *Bacillus* metabolites with established roles in antimicrobial activity, including fengycin (86% similarity), bacillaene (100%), bacilysin (100%), bacillibactin (100%), and the RiPP subtilosin A (100%).

**Table 2:**
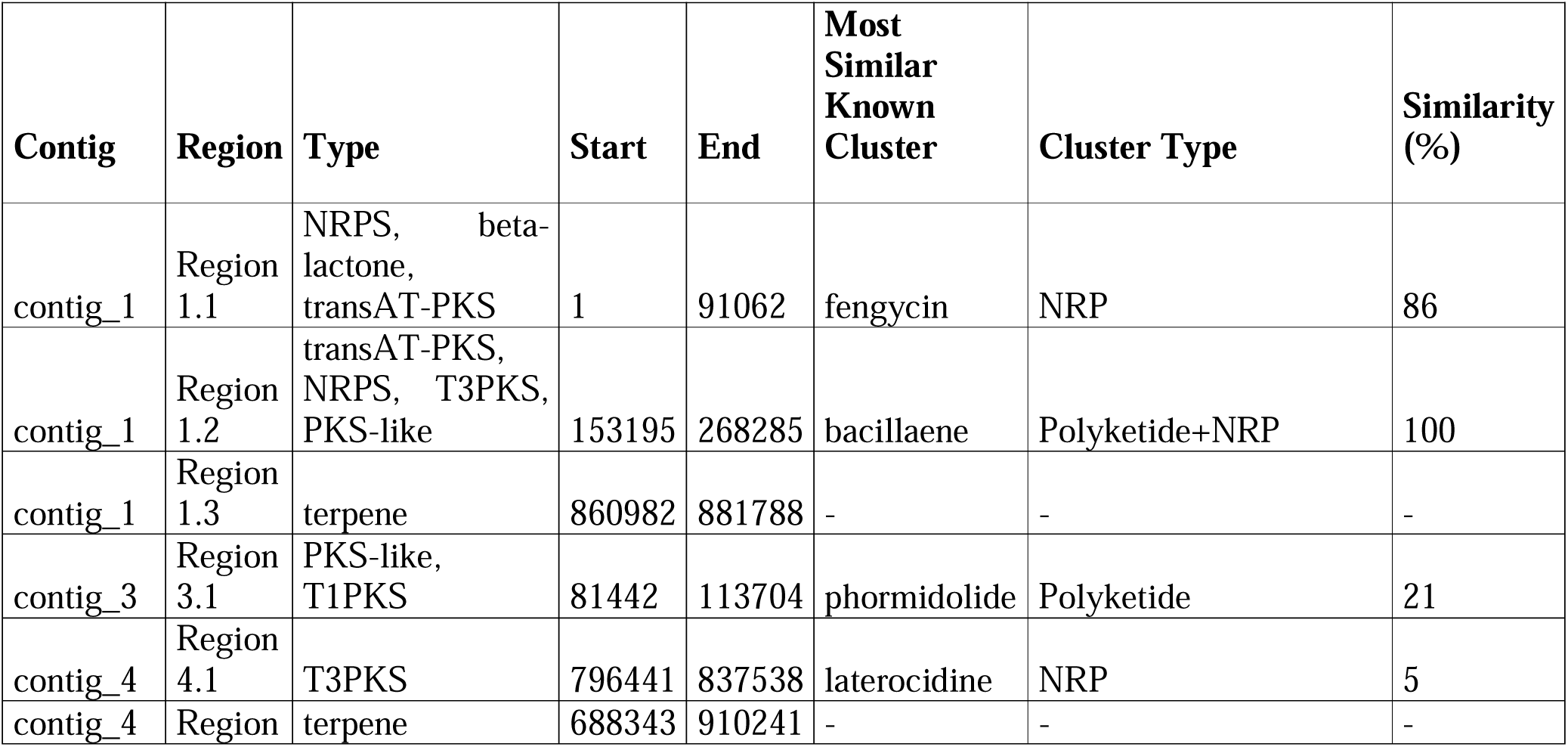
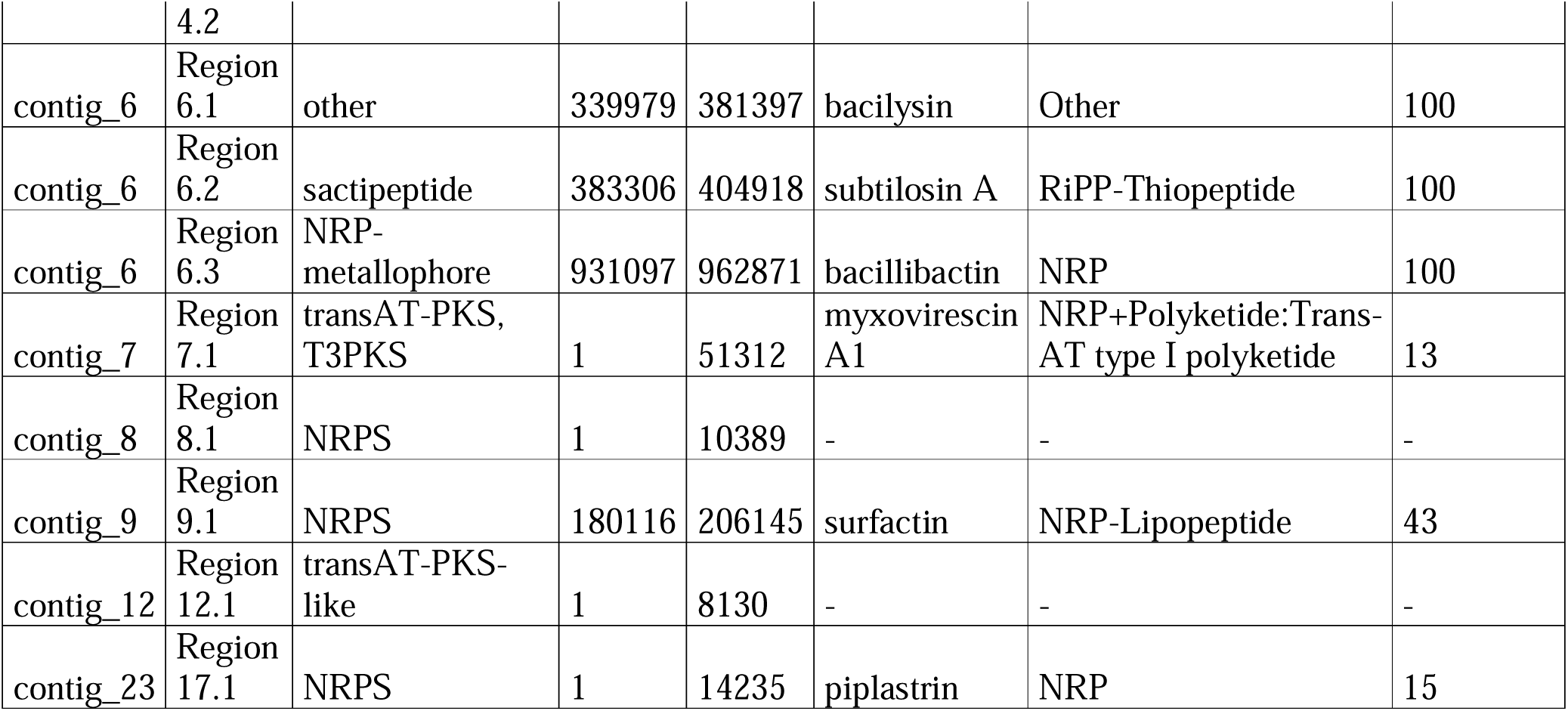
Secondary metabolite biosynthetic gene clusters predicted in the genome of the bacterial isolate using antiSMASH. . The table lists the contig, region, cluster type, genomic coordinates, the most similar known cluster, compound class, and percentage similarity. This table represents secondary metabolites producing biosynthetic clusters of DDC20 with homology to the most similar known biosynthetic clusters as predicted by antiSMASH.

Importantly, the presence of fengycin- and surfactin-like NRPS clusters is consistent with the lipopeptides detected by LC–MS, supporting their active production by DDC20. In addition to these conserved clusters, antiSMASH predicted a surfactin-like NRPS cluster with moderate similarity (43%), suggesting the possible production of structurally divergent surfactin homologues. Several additional BGCs exhibited low similarity to known reference clusters, including myxovirescin-like (13%), phormidolide-like (21%), laterocidine-like (5%), and piplastrin-like (15%) clusters. Given their low similarity scores, these predictions should be interpreted with caution and may reflect either highly divergent biosynthetic pathways or novel secondary metabolites.

The detection of multiple trans-AT PKS, NRPS, and RiPP gene clusters highlights the genetic capacity of DDC20 to produce a chemically diverse array of bioactive compounds. While only lipopeptides such as surfactins and fengycins were experimentally validated in this study, the presence of additional low-similarity BGCs suggests unexplored biosynthetic potential that warrants further functional and chemical characterization. This genomic analysis of DDC20 using antiSMASH reveals a rich secondary metabolite biosynthetic potential, particularly in the production of lipopeptides, polyketides, and RiPPs. These findings suggest that DDC20 and DDC_NEW2 could be a valuable biocontrol agent against agricultural pathogens and a potential source of novel antimicrobial compounds.

## 4. Discussion

In this study, we screened 436 bacterial strains isolated from healthy banana roots for their inhibition of Foc, and the developments of Panama disease. From these, 123 strains demonstrated in-vitro antagonistic activity against Foc, with 93 showing statistically significant inhibition. Notably, their inhibitory spectrum extended also to another F. oxysporum strain pathogenic to cannabis (FC), while only moderate activity was observed against Rhizoctonia solani (RH), indicating possible specificity within their antifungal mechanisms. To validate these findings under more accurate conditions, 64 strains of these, showing the strongest in-vitro activity, were tested in-planta in greenhouse experiments under controlled environment. We found that 22 of these strains significantly reduced disease severity in banana plants inoculated with Foc. In addition, we have examined the antifungal potential of extracellular metabolites secreted by 78 of these strains through kinetic assays using GFP-expressing Foc spores. A total of 48 exhibiting statistically significant inhibitory effects based on fluorescence derived AUDPC values.

Based on all the above screening assays, two strains, DDC20 and DDC_NEW2, consistently displayed strong antifungal activity across the multiple experiments, positioning them as best candidates for further investigation.

In-vitro assays revealed that both bacteria significantly reduced Foc mycelial growth compared to non-inhibitory controls (p ≤ 0.0002). In in-planta assay, again, DDC_NEW2 and DDC20 emerged as highly promising candidates, reducing disease severity by 60 to 67% (respectively) compared to the control without addition of inhibiting bacteria. The consistency of their in-planta effect across experimental systems and conditions, as indicated by non-significant interaction terms, further highlights their robustness and reliability in disease suppressiveness. Additionally, supernatant collected from DDC20 consistently exhibited a strong suppressive effect on Foc GFP fluorescence (p ≤ 0.003), suggesting the involvement of secreted antifungal compounds. In contrast, supernatant of DDC_NEW2 showed more variable results, with non-significant inhibition under the same conditions (p = 0.17), implying that its biocontrol activity may depend on direct bacterial-fungal interaction or in-planta mechanisms rather than diffusible metabolites alone. Taken together, these findings offer a multidimensional view of biocontrol efficacy, combining in-vitro inhibition, greenhouse disease suppression, and mechanistic insights through extracellular metabolite analysis.

Phylogenetic analysis revealed that DDC_NEW2 and DDC20 cluster within distinct Bacillus. The DDC_NEW2 strain is positioned closely to B. velezensis ZB42 (the reference bacterium in NCBI). The B. velezensis group, members of which are known for their plant growth-promoting and biocontrol properties [37, 38]. DDC20 is placed in a different subclade, clustering within the B. halotolerans group, many members of which are known for their plant growth and biocontrol properties [39–41]. These findings agree with previous studies reporting similar levels of inhibition by related Bacillus species against different phytopathogens. For instance, B. velezensis BVE7 inhibited F. oxysporum mycelial growth by 61% under in-vitro conditions [42], while B. velezensis strain SDTB038 filtrates achieved 45–66% inhibition and reduced tomato crown rot severity under greenhouse assays [43]. Comparable results have also been reported for Bacillus isolates controlling Ralstonia solanacearum and F. oxysporum in various hosts [38, 44].

To better understand the molecular mechanisms underlying the antifungal activity of the two isolates DDC20 and DDC_NEW2 observed in vitro and in planta, whole-genome sequencing was performed, followed by secondary metabolite biosynthetic gene cluster analysis using the antiSMASH platform v7.1.0 [45]. This in silico approach enabled the prediction of secondary metabolite biosynthetic gene clusters (BGCs) within the genomes, providing insight into their biosynthetic potential to produce bioactive compounds. Both strains exhibited a diverse repertoire of BGCs, including clusters encoding non-ribosomal peptide synthetases (NRPS), polyketide synthases (PKS), and ribosomally synthesized and post-translationally modified peptides (RiPPs). Notably, NRPS-derived lipopeptides such as surfactin [46–48] and fengycin [49–51], known for their potent antifungal properties, were identified in both genomes. In addition, DDC_NEW2 and DDC20 carried conserved BGCs associated with the production of siderophore bacillibactin [52], bacillaene [53, 54], and macrolactin [55], compounds with well-documented antimicrobial and antifungal activity and relevance in plant–microbe interactions. Additionally, genes coding for metalloproteases, glucanases, and other polysaccharide-degrading enzymes, predicted from the COG functional categories of DDC20 and DDC_NEW2, may complement the secondary metabolite-based defense by enzymatically targeting the fungal cell wall. Such enzymatic degradation likely weakens the structural integrity of the hyphae, facilitating subsequent membrane disruption by lipopeptides. Similar synergistic interactions between hydrolytic enzymes and lipopeptides have been well documented in Bacillus-based biocontrol systems. In particular, β-1,3-glucanases, chitinases, and proteases produced by strains such as B. velezensis and B. amyloliquefaciens facilitate the degradation of the fungal cell wall, thereby enhancing the ability of fengycins and surfactins to access and disrupt the plasma membrane of Fusarium spp [56, 57]. Similarly, metalloproteases of the Bacillolysin family have been implicated in the degradation of extracellular proteins and the destabilization of pathogen structures [58], supporting their presumed role in DDC_NEW2 and DDC20. Interestingly, several predicted biosynthetic gene clusters showed low similarity (<50%) to entries in the MIBiG database, indicating the presence of biosynthetic pathways with limited homology to currently characterized clusters. While the chemical products and biological functions of these clusters remain unknown, their genomic organization suggests potential biosynthetic diversity beyond well-characterized Bacillus metabolites. However, a limitation of the antiSMASH-based analysis is that several clusters were located at contig boundaries, raising the possibility of assembly-related truncation. Such incomplete assemblies, which are common in short-read sequencing data, may lead to partial or inaccurate annotation of biosynthetic pathways, particularly for large or hybrid PKS–NRPS clusters whose functional domains can span contig edges. Future functional characterization will be required to determine whether any of these low-similarity clusters contribute to the antifungal activity observed in-vitro and in-planta.

These bioinformatic predictions aligned well with the LC-MS data obtained from bacterial supernatants. The profiles of strains DDC20 and DDC_NEW2 confirmed the production of cyclic lipopeptides, including surfactins (m/z 1008–1036) and fengycins (m/z 1449–1505), identified based on their retention times and molecular weights. These compounds were previously reported in multiple Bacillus species with strong antifungal activity [59–62], disrupting fungal membranes, inducing leakage of intracellular contents, and inhibiting spore germination [63–65] . DDC20 exhibited a broader and more complex metabolite pattern, including several unidentified peaks absent in DDC_NEW2, suggesting the synthesis of additional and/or structurally novel secondary metabolites. These findings correlate well with the antiSMASH genomic predictions, which identified numerous BGCs in both strains, including those encoding surfactin, fengycin, bacillaene, and mojavensin, as well as multiple orphan NRPS–PKS clusters of low similarity to known references. The clear verification of the genomic by the metabolomic data underscores the robust anti-fungal potential of both isolates and reinforces the integrative strength of combining silico and analytical approaches in microbial biocontrol research [66, 67].

The identified amphiphilic molecules such as surfactins are known to reduce surface tension, form membrane pores, and induce cell leakage, mechanisms that align well with the hyphal deformation and chlamydospore formation observed under microscopic analysis [68–70]. In fact, our microscopic observations provided direct visual evidence supporting the antifungal activity of DDC20 and DDC_NEW2. Co-cultivation of Foc with either DDC20 or DDC_NEW2 led to marked hyphal deformation, including swelling, irregular branching, cytoplasmic granulation, and fragmentation, as well as abundant chlamydospores, all typical of fungal stress and survival responses under hostile conditions. These morphological alterations are consistent with cell-wall weakening and membrane destabilization induced by antimicrobial metabolites or hydrolytic enzymes, phenomena that have also been documented in other Bacillus–Fusarium interactions [71, 72]. This is supported by the PI staining of bacterial CFCM treated Foc since PI can only penetrate damaged membrane cells. This evidence of membranal damage was especially pronounced for DDC20 treatments, in which extensive hyphal disintegration and cytoplasmic leakage were observed, revealing severe membrane perturbation. Such results mirror previous reports in which Bacillus-produced lipopeptides (notably surfactin and fengycin) and proteolytic enzymes synergistically disrupt fungal membranes and induce cell death [59, 60]. Similar morphological and viability disruptions have been reported in B. velezensis and B. amyloliquefaciens strains acting against Fusarium spp., R. solani, and Botrytis cinerea [42, 73, 74], confirming that the phenomena observed here align with the established biocontrol repertoire of other Bacillus.

Conversely, fengycins, which exhibit high specificity against filamentous fungi by targeting membrane sterols, were detected at lower amount consistent with their more polar nature and reduced extractability under the tested conditions. This polarity-dependent extraction was further confirmed by solvent comparisons: butanol (BuOH), being more polar than ethyl acetate (EA), yielded higher amounts of fengycins and other unidentified polar metabolites, whereas EA preferentially recovered the less polar surfactins. DDC20, which exhibited stronger antifungal activity in both in-vitro and in-planta assays, produced a higher proportion of polar, unidentified metabolites that were efficiently extracted with BuOH. This observation raises the possibility that some of these unknown polar compounds contribute significantly to their superior biocontrol efficacy. Similar polarity-driven recovery trends have been reported in B. velezensis and B. amyloliquefaciens, where solvent extraction significantly influences metabolite composition and apparent antifungal potency [75, 76].

Interestingly, the metabolite profiles of both DDC20 and DDC_NEW2 remained unchanged, without elicitation, in the presence of live or dead Foc spores, indicating that antifungal compound production is constitutive rather than inducible. Such constitutive secretion of lipopeptides ensures a continuous defensive barrier within the rhizosphere, a trait advantageous for sustained disease suppression under natural conditions [77–79].

The fractionation experiments provided additional insight into the chemical basis underlying the superior antifungal activity of strain DDC20 compared with DDC_NEW2. While all crude extracts reduced Foc GFP fluorescence relative to the untreated control, clear strain-dependent differences were observed in both the magnitude and robustness of inhibition. Extracts derived from DDC_NEW2 (fractions A–D) displayed moderate antifungal activity, with butanol fractions generally outperforming ethyl acetate fractions, consistent with the enrichment of polar lipopeptides such as fengycins in the BuOH phase. However, the overall inhibitory capacity of DDC_NEW2 extracts remained limited compared to those of DDC20. Importantly, the fractionation data does not support fungal elicitation of secondary metabolite production by DDC_NEW2 and DDC20. Cultures grown in the presence of live or dead fungal material yielded comparable LC–MS profiles and similar overall antifungal potency, indicating that metabolite biosynthesis in DDC_NEW2 and DDC20 is constitutive rather than induced by fungal signals. The minor differences observed between certain extract combinations therefore reflect interactions between chemically distinct metabolite pools rather than differential biosynthesis triggered by fungal exposure.

In DDC_NEW2, specific interaction patterns between fractions were observed. However, LC–MS analyses demonstrated that secondary metabolite production was constitutive and not influenced by the presence of either live or dead fungal material. The combination of ethyl acetate and butanol fractions resulted in stronger inhibition than either fraction alone, suggesting a synergistic interaction between apolar and polar metabolites. In contrast, all fractions derived from DDC20 (E–H) exhibited markedly stronger antifungal activity than those of DDC_NEW2, with inhibition approaching near-complete suppression of GFP fluorescence. Because DDC20 extracts already displayed maximal or near-maximal inhibition, potential synergistic or additive effects between fractions could not be resolved under the conditions tested. LC–MS analyses indicate that DDC20 produces not only surfactins and fengycins, but also additional polar metabolites absent or less abundant in DDC_NEW2, which likely contribute collectively to its enhanced antifungal efficacy.

Together, these fractionation results support the finding that both bacteria secrete highly active metabolites and that DDC20 secretes a broader and more potent chemical arsenal than DDC_NEW2, combining major lipopeptide families with additional, unidentified metabolites. Similar findings have been reported for B. velezensis and B. amyloliquefaciens, where high-performing strains exhibit enhanced antifungal activity due to the combined action of multiple lipopeptides and accessory metabolites rather than a single compound class [68, 80]. Our results align with this model, demonstrating that these highly effective Bacillus biocontrol strains rely on chemically diverse and constitutively produced antifungal arsenal capable of targeting multiple fungal physiological processes.

From an application perspective, the constitutive production of antifungal metabolites by DDC20 and DDC_NEW2 suggests that these strains could be deployed either as live biocontrol inoculants or as sources of bioactive extracts. While live bacterial application would allow rhizosphere colonization and continuous metabolite production, the strong activity observed for cell-free supernatants and crude extracts (particularly from DDC20) also supports the potential use of extracted or even industrially synthesized metabolites as separate antifungal formulations or in combination with live bacteria, depending on agronomic and regulatory constraints.

## 5. Conclusion

Our integrative pipeline, covering in-vitro assays, greenhouse experiments, microscopy, genome analysis, and LC–MS metabolite profiling demonstrated that the isolates are potent biocontrol candidates against F. oxysporum f. sp. cubense (TR4). Both strains consistently reduce Foc growth and disease, with DDC20 showing superior efficacy. Mechanistically, antiSMASH predictions and metabolite profiles converge on constitutive production of antifungal lipopeptides (notably surfactins and fengycins), complemented by an enzymatic arsenal inferred from genomic functions; confocal and light microscopy corroborate membrane damage and hyphal dysfunction. LC–MS indicates that DDC20 has broader, more polar metabolite repertoire than DDC_NEW2, suggesting additional, yet-uncharacterized antifungal molecules. Together, these findings support DDC20 (and to a lesser extent DDC_NEW2) as scalable, environmentally friendly BCAs for Panama disease management and motivate targeted purification and characterization of the unidentified metabolites, alongside transcriptomic/field validation to optimize deployment in banana production systems.

