## Supplementary material for "Isolation and characterization of novel banana rhizosphere bacteria for the control of *Fusarium oxysporum* f. sp. *cubense* TR4": Fig. S

### Slide 1
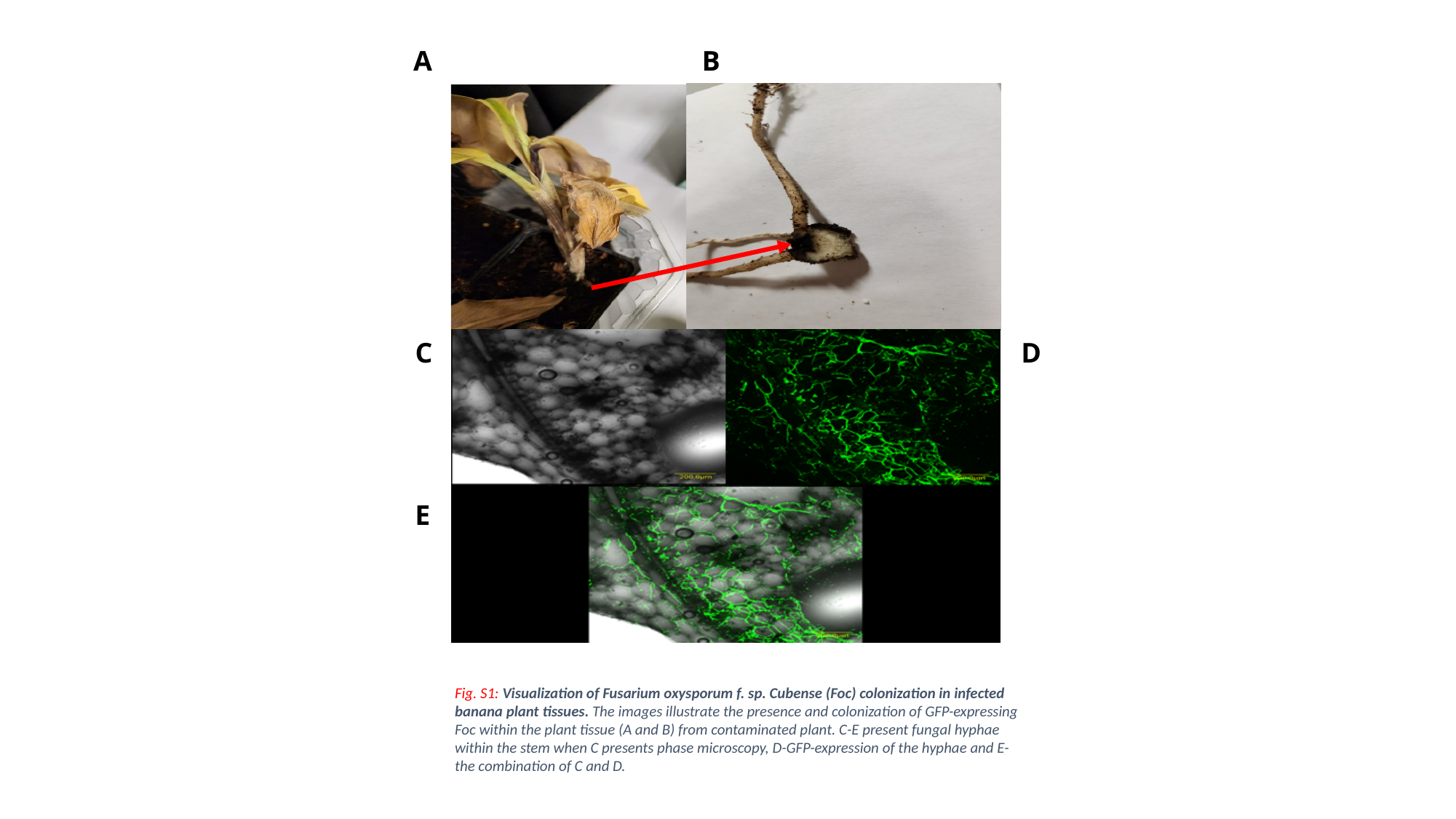

A
B
Fig. S1: Visualization of Fusarium oxysporum f. sp. Cubense (Foc) colonization in infected banana plant tissues. The images illustrate the presence and colonization of GFP-expressing Foc within the plant tissue (A and B) from contaminated plant. C-E present fungal hyphae within the stem when C presents phase microscopy, D-GFP-expression of the hyphae and E-the combination of C and D.
C
D
E

### Slide 2
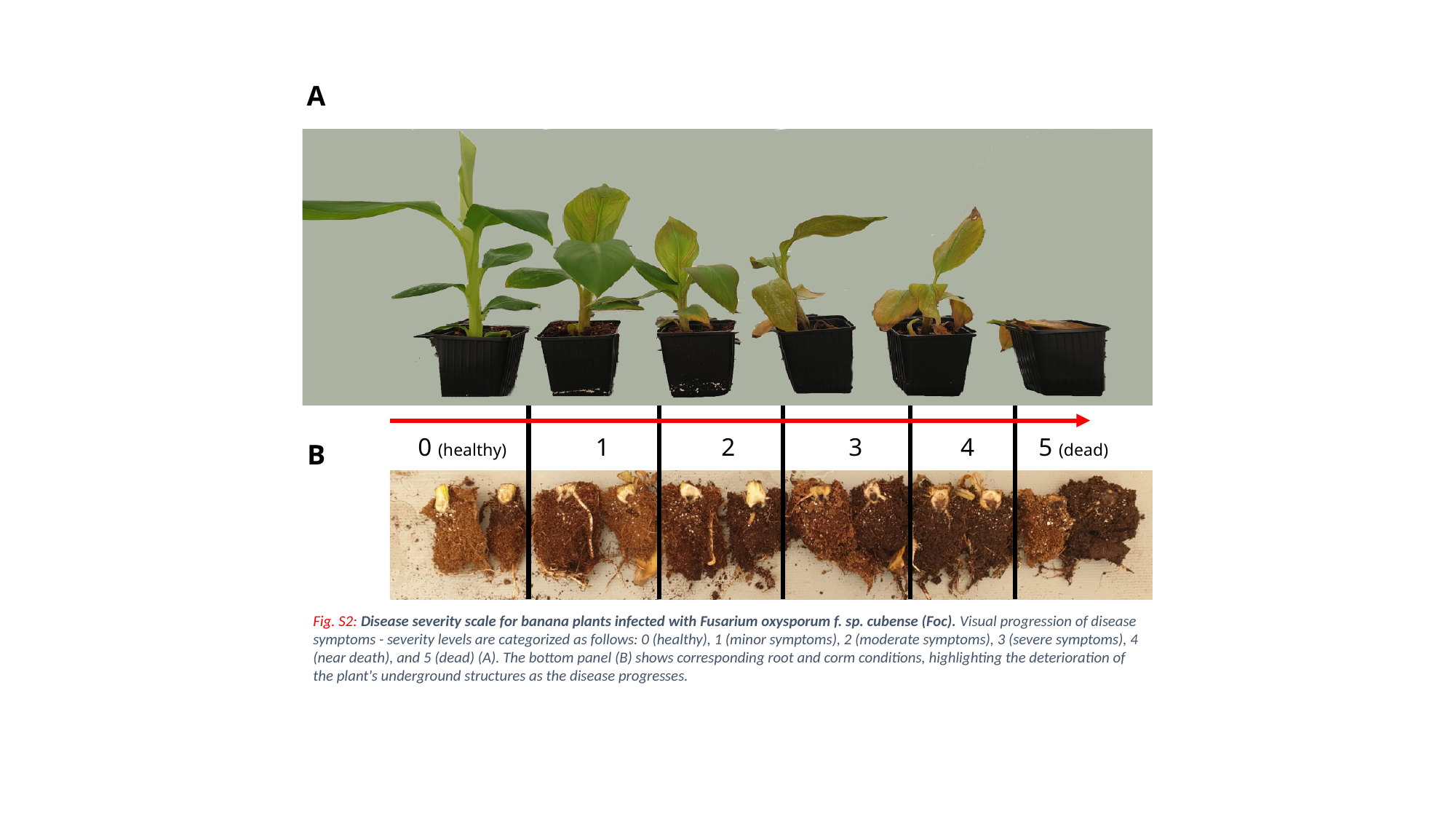

A
0 (healthy)
1
2
3
4
5 (dead)
Fig. S2: Disease severity scale for banana plants infected with Fusarium oxysporum f. sp. cubense (Foc). Visual progression of disease symptoms - severity levels are categorized as follows: 0 (healthy), 1 (minor symptoms), 2 (moderate symptoms), 3 (severe symptoms), 4 (near death), and 5 (dead) (A). The bottom panel (B) shows corresponding root and corm conditions, highlighting the deterioration of the plant's underground structures as the disease progresses.
B

### Slide 3
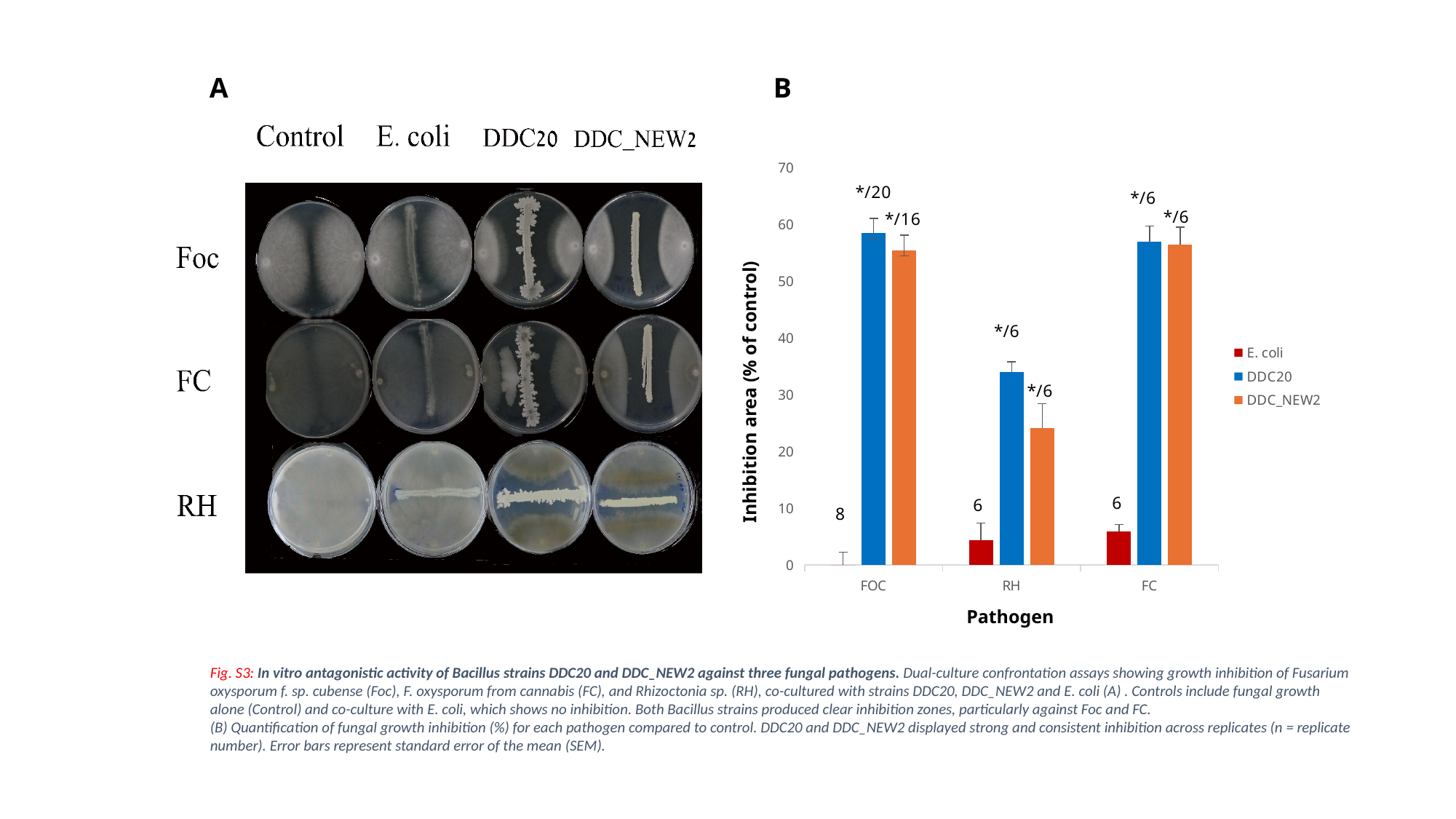

A
B
#### Chart
| Category | E. coli | DDC20 | DDC_NEW2 |
|---|---|---|---|
| FOC | 0.0 | 58.53010213129974 | 55.455000678093405 |
| RH | 4.334307484954025 | 33.954521798310566 | 24.048200488410252 |
| FC | 5.9098732081390715 | 56.90492788985229 | 56.377173107925586 |Inhibition area (% of control)
Pathogen
Fig. S3: In vitro antagonistic activity of Bacillus strains DDC20 and DDC_NEW2 against three fungal pathogens. Dual-culture confrontation assays showing growth inhibition of Fusarium oxysporum f. sp. cubense (Foc), F. oxysporum from cannabis (FC), and Rhizoctonia sp. (RH), co-cultured with strains DDC20, DDC_NEW2 and E. coli (A) . Controls include fungal growth alone (Control) and co-culture with E. coli, which shows no inhibition. Both Bacillus strains produced clear inhibition zones, particularly against Foc and FC.
(B) Quantification of fungal growth inhibition (%) for each pathogen compared to control. DDC20 and DDC_NEW2 displayed strong and consistent inhibition across replicates (n = replicate number). Error bars represent standard error of the mean (SEM).

### Slide 4
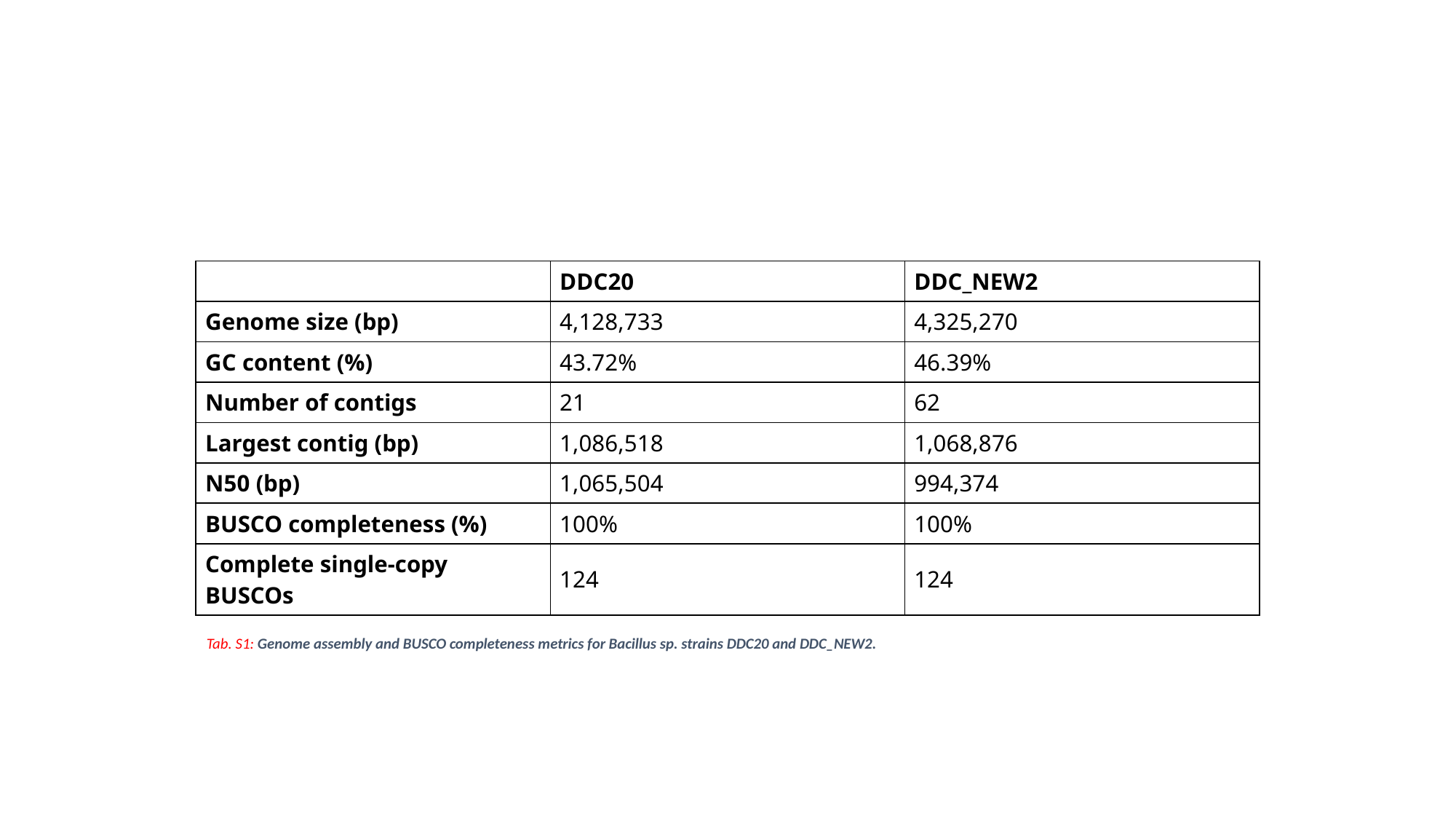

| | DDC20 | DDC\_NEW2 |
| --- | --- | --- |
| Genome size (bp) | 4,128,733 | 4,325,270 |
| GC content (%) | 43.72% | 46.39% |
| Number of contigs | 21 | 62 |
| Largest contig (bp) | 1,086,518 | 1,068,876 |
| N50 (bp) | 1,065,504 | 994,374 |
| BUSCO completeness (%) | 100% | 100% |
| Complete single-copy BUSCOs | 124 | 124 |
Tab. S1: Genome assembly and BUSCO completeness metrics for Bacillus sp. strains DDC20 and DDC_NEW2.

### Slide 5
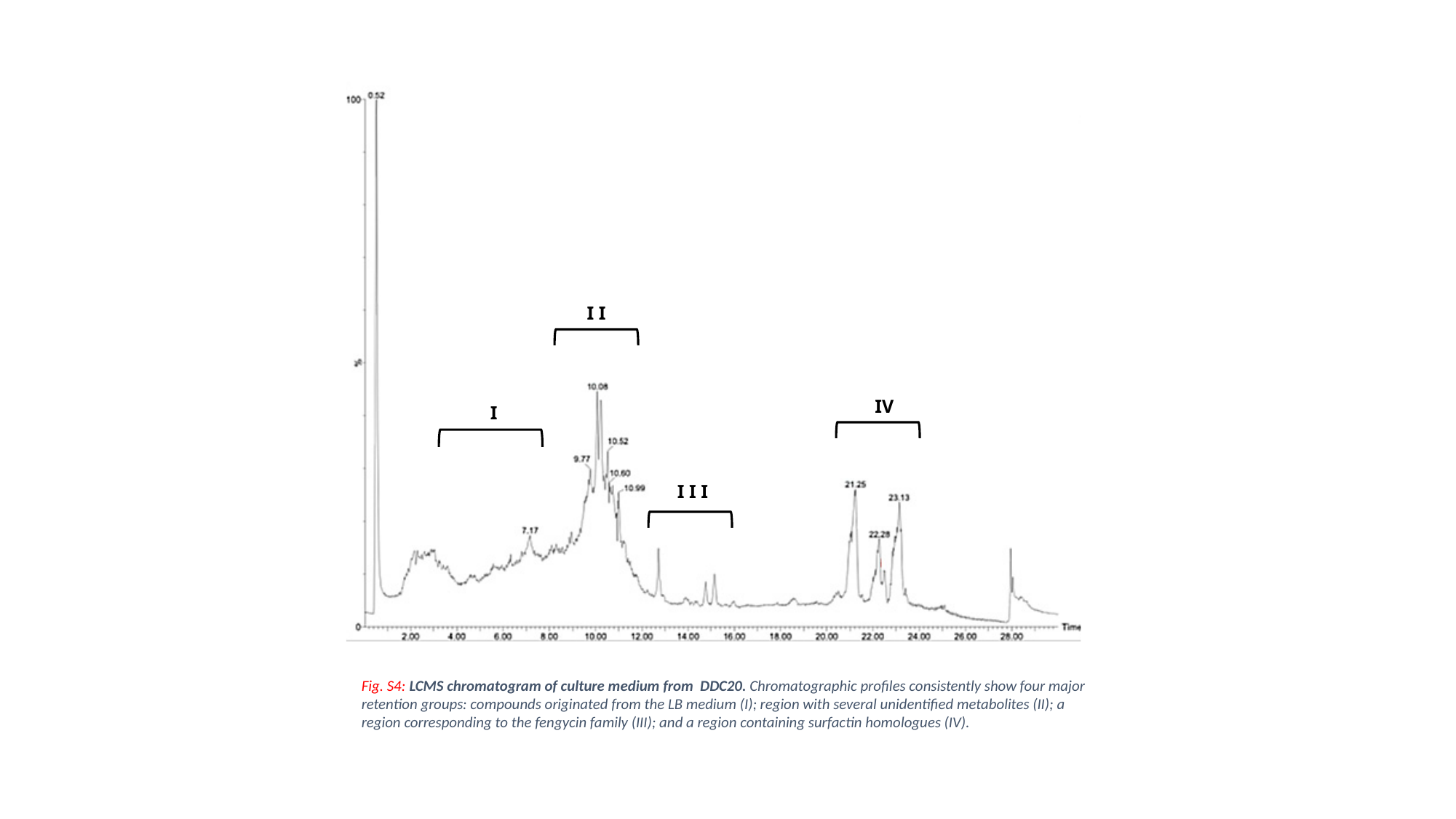

Fig. S4: LCMS chromatogram of culture medium from DDC20. Chromatographic profiles consistently show four major retention groups: compounds originated from the LB medium (I); region with several unidentified metabolites (II); a region corresponding to the fengycin family (III); and a region containing surfactin homologues (IV).
I I
IV
I
I I I

### Slide 6
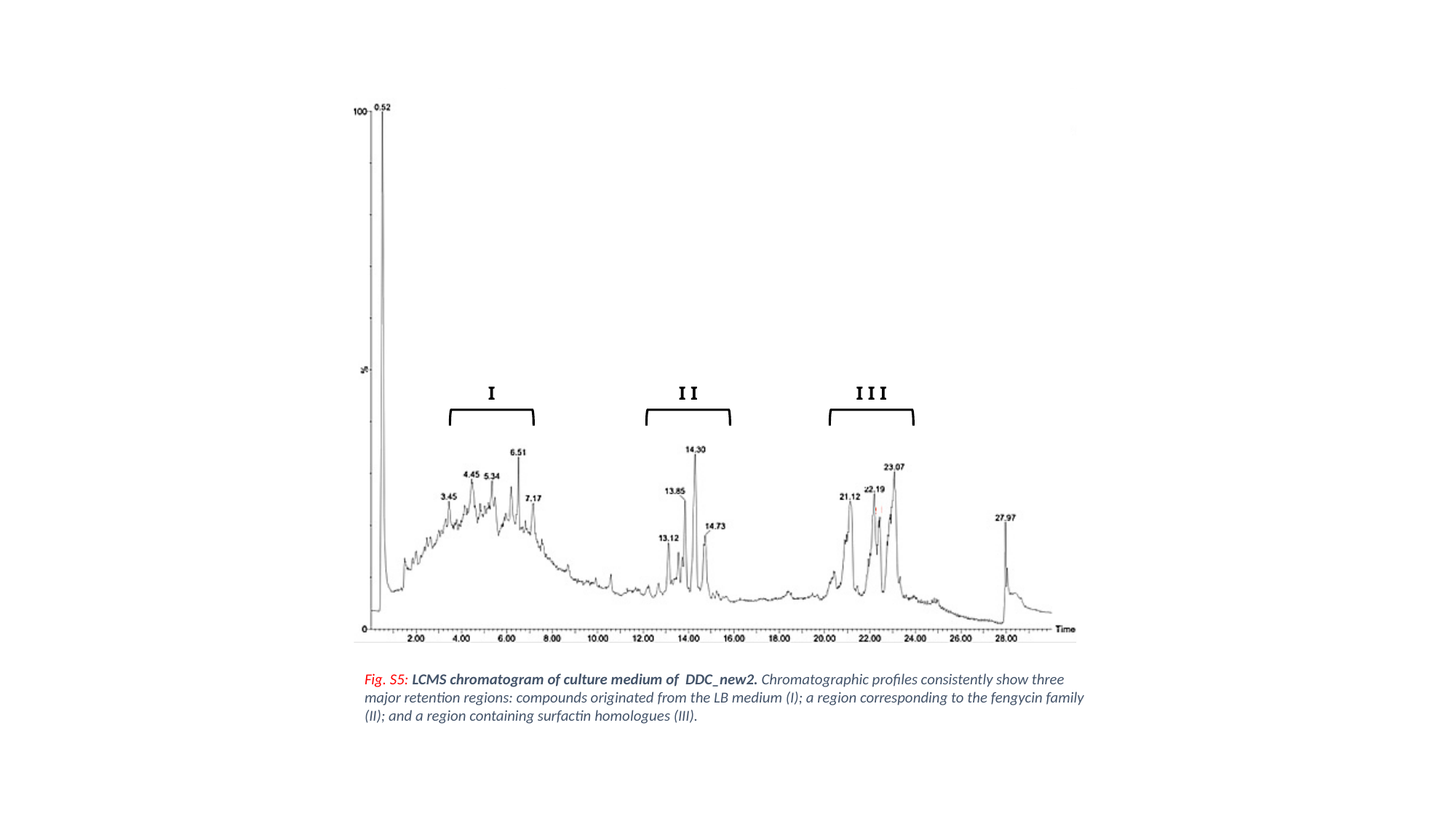

I I
I I I
I
Fig. S5: LCMS chromatogram of culture medium of DDC_new2. Chromatographic profiles consistently show three major retention regions: compounds originated from the LB medium (I); a region corresponding to the fengycin family (II); and a region containing surfactin homologues (III).
